# PWWP-ADD and N-terminal domains of DNMT3B1 confer specificity for developmentally regulated CpG island methylation

**DOI:** 10.1101/2024.10.22.619628

**Authors:** Komal Yasmin, Tatyana B Nesterova, Neil Brockdorff

**Author notes:** Babraham Institute, Babraham Research Campus, Cambridge, CB22 3AT (Komal Yasmin).

## Abstract

CpG methylation in mammalian genomes is established by the closely related de novo DNA methyltransferases DNMT3A and DNMT3B. Whilst both enzymes contribute to pervasive genome-wide CpG methylation, DNMT3B has a unique role in developmentally regulated CpG island (CGI) methylation both on the inactive X chromosome and at other sites in the genome. The mechanistic basis for this specificity is poorly understood. Here we have developed an in vitro embryonic stem cell model system to dissect critical determinants of DNMT3B specificity. Our model faithfully recapitulates developmentally regulated CGI methylation and additionally provides novel insights into CpG methylation at cis-regulatory elements. Using genetic complementation, we show that DNMT3B specificity is attributable solely to the catalytic isoform DNMT3B1. Domain swap experiments demonstrate an important role both for the PWWP-ADD chromatin binding and unstructured N-terminal domains. Together, these findings advance our mechanistic understanding of the unique roles of DNMT enzymes in establishing CpG methylation in development.

## Introduction

In vertebrate genomes 5-methyl cytosine (5meC) occurs pervasively at CpG dinucleotides where it is important for regulating both the activity of transposable elements and expression of developmentally regulated genes (reviewed in Greenberg & Bourc’his, 2019; Schübeler, 2015). CpG methylation is catalysed by DNA methyl transferases (DNMTs) which can be classified into two categories based on their DNA substrate preference. First, *de novo* methyltransferases (including in mammals DNMT3A and DNMT3B) which establish methylation of previously unmethylated CpG sites on DNA and second, the maintenance methyltransferase (DNMT1) which propagates CpG methylation from hemi-methylated DNA on the parent strand to the daughter strand during DNA replication (reviewed in (Gowher & Jeltsch, 2018; Schübeler, 2015).

Whilst vertebrate CpG methylation is largely pervasive, an important exception occurs at CpG islands (CGIs), elements typically 1-2kb in length located at the 5’ end of a large proportion of genes (Bird, Taggart, Frommer, Miller, & Macleod, 1985). Despite having a higher than average density of CpG sites, in most instances CGIs remain unmethylated throughout development, regardless of whether the associated genes are expressed (reviewed in Illingworth & Bird, 2009). The basis for CGI hypomethylation is not fully understood but evidence points to at least two default pathways. First, the histone modification H3K4me3, which is enriched at CGIs acts to inhibit the activity of the vertebrate de novo DNA methyltransferases, DNMT3A and DNMT3B (Guo et al., 2015). H3K4me3 deposition at CGI is at least in part attributable to the presence of a CXXC domains in specific H3K4 histone methyltransferases which recognises unmethylated CpG dinucleotides (Birke et al., 2002; Cierpicki et al., 2010; Long, Blackledge, & Klose, 2013). Secondly, CXXC recognition of CGIs by the enzymes TET1 and TET2 that catalyse removal of cytosine methylation via a hydroxymethylated intermediate (Lorsbach et al., 2003; Ono et al., 2002). Loss of function of TET1 and TET2 is linked to a gain of methylation at CGIs (Dawlaty et al., 2013; Ficz et al., 2011; Hon et al., 2014; Ito et al., 2010; Wu et al., 2011).

Although the majority of CGIs remain hypomethylated throughout development, there are circumstances in which CGI methylation does occur (Auclair, Guibert, Bender, & Weber, 2014; Borgel et al., 2010; Dahlet et al., 2020; Straussman et al., 2009). An example of this is CGIs associated with meiosis and germline-specific genes that in somatic cells are silenced by a repressor complex that includes the PRC1 polycomb protein PCGF6 (Borgel et al., 2010; Dahlet et al., 2020; Mitsuhiro et al., 2017). In this instance CGI methylation is largely dependent on the de novo DNA methyltransferase DNMT3B, rather than its closely related and broadly co-expressed homologue, DNMT3A (Auclair et al., 2014; Borgel et al., 2010). A second example of developmentally regulated CGI methylation is during X chromosome inactivation (XCI) in the cells of female mammals. Here, CGI on the inactive X chromosome (Xi) gain CpG methylation, also in a DNMT3B-dependent manner (Gendrel et al., 2012). CGI methylation is important for the maintenance of XCI, occurring downstream of initiation and establishment of chromosome silencing by the cis-acting non-coding RNA Xist (Lock, Takagi, & Martin, 1987; Sado et al., 2000). Prior studies have established that methylation of the majority of Xi CGIs is additionally dependent on the chromosomal protein SmcHD1 which is recruited to the Xi at a relatively late developmental stage, 2-3 days after the onset of cellular differentiation (Bowness et al., 2022; Gendrel et al., 2012). A final example where CGI methylation is observed is at tumour suppressor genes (TSG) in cancer cells. Little is known about specific mechanisms in this instance although stochastic gain of methylation and accompanying selection for TSG repression during cancer evolution are likely important determinants (reviewed in (Baylin & Jones, 2016).

DNMT3A and DNMT3B are highly homologous with around 85% amino acid similarity. Protein architecture is also highly similar with a C-terminal catalytic domain and centrally located PWWP and ADD domains, both with ascribed roles in chromatin recognition. The N-terminal regions are relatively diverse and are predicted to be unstructured (reviewed in (Jeltsch & Jurkowska, 2016). Two recent studies have reported that the N-terminal region of DNMT3A can bind to the histone modification H2AK119ub1 which is catalysed by PRC1 Polycomb complexes (Gu et al., 2022; Weinberg et al., 2021). Consistent with similarities in the features of DNMT3A and DNMT3B, gene knockouts in ES cells indicate that the two proteins are in large part redundant in their role in establishing de novo CpG methylation patterns. Gene knockout results in mid-late embryo lethality in both cases (Okano, Bell, Haber, & Li, 1999). Despite these overall similarities DNMT3A and DNMT3B do have more specific functions, including, as above, in CGI methylation. The mechanistic basis of these specificities, including the role of DNMT3B in CGI methylation is not well understood. A priori, specific functions could be attributable to differences in protein structure and/or the presence of non-catalytic isoforms, notably DNMT3L and DNMT3B3, both of which can modulate DNMT3A/B catalytic activity through the formation of heterodimers (Chedin, Lieber, & Hsieh, 2002; Chen, Mann, Hsieh, Riggs, & Chédin, 2005; Gordon, Hartono, & Chédin, 2013; Gowher, Liebert, Hermann, Xu, & Jeltsch, 2005; Suetake, Shinozaki, Miyagawa, Takeshima, & Tajima, 2004).

In this study we set out to better define DNMT3B specificity in CGI methylation. Accordingly, we established an ES cell-based model that recapitulates developmentally regulated CGI methylation both on Xi and genome wide. By engineering defined perturbations to modify DNMT3B we demonstrate that specificity is conferred by DNMT3B1, the catalytic isoform of the protein with critical roles for the PWWP-ADD domain and the unstructured N-terminal region.

## Results

### Establishment of an ES cell model that recapitulates Xi and developmentally regulated CGI methylation

We set out to develop an ES cell-based model to dissect the mechanisms governing specificity of DNMT3B for Xi and developmentally regulated CGI methylation. Accordingly, we made use of iXist-ChrX, an interspecific XX female mESC line in which the native Xist promoter on one allele is replaced with the TetOn promoter for Doxycycline inducible expression (Nesterova et al., 2019). To test the suitability of this experimental system to study CGI methylation during XCI, we performed MBD-seq at various timepoints during embryoid body (EB) differentiation in the presence of doxycycline and LIF withdrawal (Figure 1A, B). MBD-seq employs the MBD domain of MBD1 for affinity purification of methylated CpG-dense DNA. To correct for variability in IP efficacy or library preparation between experiments, we included human DNA to serve as a calibration control. Figure 1C-D shows CGI methylation during EB differentiation and Xist gene induction. As expected, we observed a proportion of CGIs across all chromosomes gain CpG methylation over the EB differentiation time course (Figure 1C), whilst on ChrX we observed CpG methylation gain on the majority (382/448) of CGIs (Figure 1D, lower panel). Selected examples of autosomal and ChrX CGI that gain CpG methylation are illustrated in Figure 1E. We also observed a significant number of non-CGI MBD peaks, lying in most cases >2 kb from the nearest transcription start site (TSS). These peaks are discussed in detail below.

**Figure 1.**
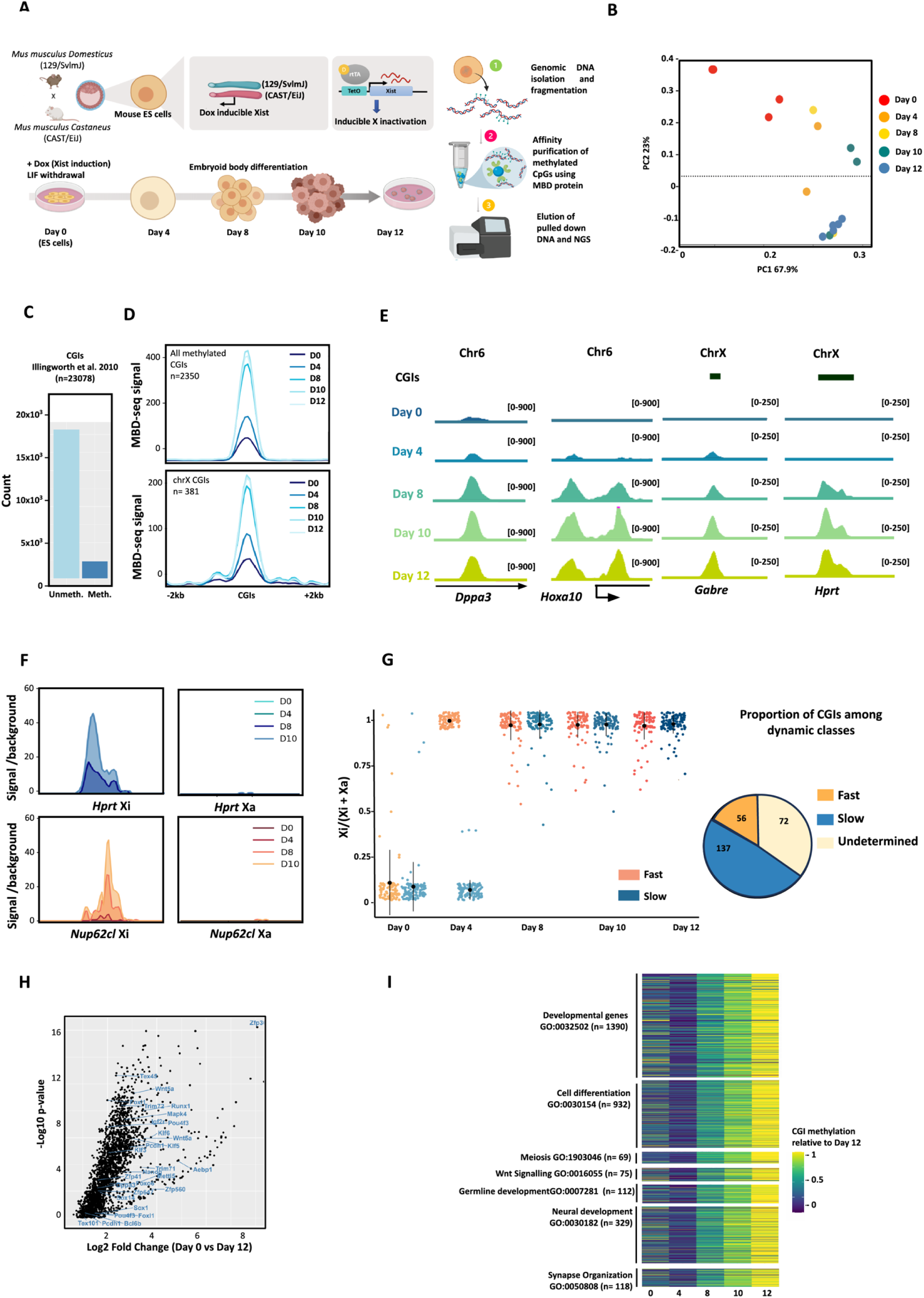
iXist-ChrX cells as a model to analyse developmentally regulated CGI methylation. A) Schematic depiction of iXist-ChrX interspecific female mouse ES cell line engineered to contain a Tet-inducible *Xist* promoter on the *M. Castaneus* derived X chromosome, subjected to EB differentiation and MBD-seq. B) PCA plot showing clustering for the MBD-seq replicates collected for day 0 – day 12 of EB differentiation. C) Barplot showing the proportion of CGIs that undergo methylation during EB differentiation in iXist-ChrX cells (defined by DeSEQ2 on MBD-seq data collected from three replicates each of day 0 ES cells and day 12 EB, FDR < 0.01). D) Representative metaplot showing the average profile of all CGIs (top) or X-linked CGIs (bottom) during EB differentiation. D0-D12 represent day 0 - day 12 of EB differentiation. E) Genome browser tracks showing gain of DNA methylation at illustrative autosomal and X-linked CGIs during EB differentiation. *Gabre* CGI exemplifies fast DNA methylation on the Xi (from day 4) and *Hprt* CGI exemplifies slow CGI methylation on the Xi. F) Metaplots for MBD-seq data at *Hprt* and *Nup62cl* CGIs after allelic split. MBD-seq signal maps largely to Xi. G) Allelic ratio (AR) analysis for X-linked CGIs shows two main trends. A small proportion (quantified in pie chart on right) of “Fast methylating” CGIs reach AR of 1 at day 4. A larger class of “slow methylating CGIs” attain AR=1 at day 8 of EB differentiation. Owing to strong bias of MBD-seq signal from Xi (panel D) the allelic ratio (Xi/(Xi+Xa)) for MBD-seq data becomes Xi/(Xi+0) = 1. Every dot represents an individual CGI on ChrX. “Undetermined CGIs” in the pie chart represent the group that could not be confidently assigned owing to low sequencing depth after allelic split. H) Volcano plot showing autosomal CGIs that are methylated during EB differentiation (expressed as log2 fold change in methylation between ES cells and EBs). Examples of CGIs associated with genes involved in developmental regulation and chromatin modification have been highlighted. The plot summarizes results from differential analysis on three replicates for MBD-seq from ES cells and 5 replicates for day 12 EBs. I) Heatmaps show the methylation of CGIs in selective GO terms during the 5-day EB timecourse. The methylation level relative to EB day 12 is shown for each CGI.

For ChrX CGIs we were able to assign MBD-seq reads allelically and this analysis confirmed gain of CpG methylation occurs largely on the Xi. Representative examples are shown in Figure 1F. For each independent CGI on the X chromosome, we defined the earliest time point during EB differentiation at which the MBD-seq signal first showed allelic bias i.e. a larger proportion of MBD-seq reads originating from Xi, (Xi/ (Xi +Xa)) > 0.5). Two trends emerged from this analysis; a small group of 56 CGIs reached an allelic ratio of 1 (AR = 1) at day 4 with levels plateauing through to day 12, referred to herein as “fast methylating” CGIs (Figure 1G). Many of the remaining CGIs acquired CpG methylation progressively reaching an allelic ratio of 1 at around 8 days of EB differentiation, referred to herein as ‘slow methylating’ CGIs (Figure 1G). This data agrees well with our previous work using non-allelic microarray-based analysis showing that Xi CGI methylation occurs in fast and slow phases, with the latter exhibiting dependence on SMCHD1 for their methylation (Gendrel et al., 2012). Finally, a small group of 20 CGIs showed gain of CGI methylation on both the Xa and Xi. Here, the number of reads from Xi + Xa increased during the time course, but the allelic ratio remained 0.5 (Figure S1A). Some CGIs in this group, are associated with germline-specific genes such as *Dusp9, Rhox13, Taf7l, AU022751,* that have been reported previously to gain CGI methylation after mouse embryo implantation (Auclair et al., 2014).

In addition to X-linked CGIs, a significant number of autosomal CGIs gained CpG methylation through the EB differentiation time course (Figure 1D, E, H). Selected examples are shown in Figure S1B. Major categories by GO analysis are developmental genes and germline-expressed genes (Figure 1I). This observation is consistent with prior studies characterising genes that gain CGI methylation during normal development (Auclair et al., 2014) and indeed we observe a close overlap of the identified gene sets (Figure 1I and Figure S1B).

We further validated our observations on CGI methylation in an alternative model, homogenous NPC (Neuronal Progenitor Cell) differentiation. MBD-seq was performed on cells collected at day 0, 12 and 15 of NPC differentiation (Figure S1C). Similar to the EB differentiation models, we were able to categorise Xi CGIs into fast- and slow-dynamic classes and the identity of CGIs in the two categories largely matched between the two differentiation systems (Figure S1C-G). The exact timing of CGI methylation onset however was relatively slow in NPC differentiation, and for this reason we used EB differentiation as the main experimental system for other experiments described below.

Together these results demonstrate that differentiating iXist-ChrX cells provide a useful model that faithfully recapitulates key features of Xi and developmental CGI methylation.

### The role of DNMT3B in CGI methylation

We went on to apply the iXist-ChrX model to analyse the specific role of DNMT3B in CGI methylation, generating a gene knockout of *Dnmt3b* using CRISPR/Cas9-mediated genome engineering (Figure 2A). *Dnmt3b^-/-^* cells behaved similarly to the wild-type (wt) cells in culture, and we observed no changes in morphology, growth rate, ability to differentiate to EBs or transcriptional silencing of the Xi chromosome (Figure S2A-C). We then performed MBD-seq on *Dnmt3b^-/-^* cells differentiated to day 12 EBs. The allelic ratio analysis revealed that CGIs on the Xi are largely unmethylated in the absence of DNMT3B (Figure 2B-C). Results for independent *Dnmt3b^-/-^*clones and replicates were fully consistent (Figure S2D,E). Since *Dnmt3b^-/-^* cells have normal levels of DNMT3A, this experiment validates that DNMT3A is not redundant with DNMT3B for methylation of the majority of CGIs on the Xi. However, we did observe that 15/265 of the Xi CGIs that are amenable to allelic analysis in wt cells gained CpG methylation biallelically in the absence of DNMT3B. Examples include *Dusp9, Kcne1l, G530011O0Rik* and *Zfp92* which are methylated independent of XCI (Figure S2F). Only 5 out of these 15 DNMT3B-independent CGIs reached an allelic ratio of 1, indicating strong allelic bias towards Xi (Figure S2F). Thus, DNMT3B is the predominant (but not the sole) de novo methyltransferase required for Xi CGI methylation.

**Figure 2.**
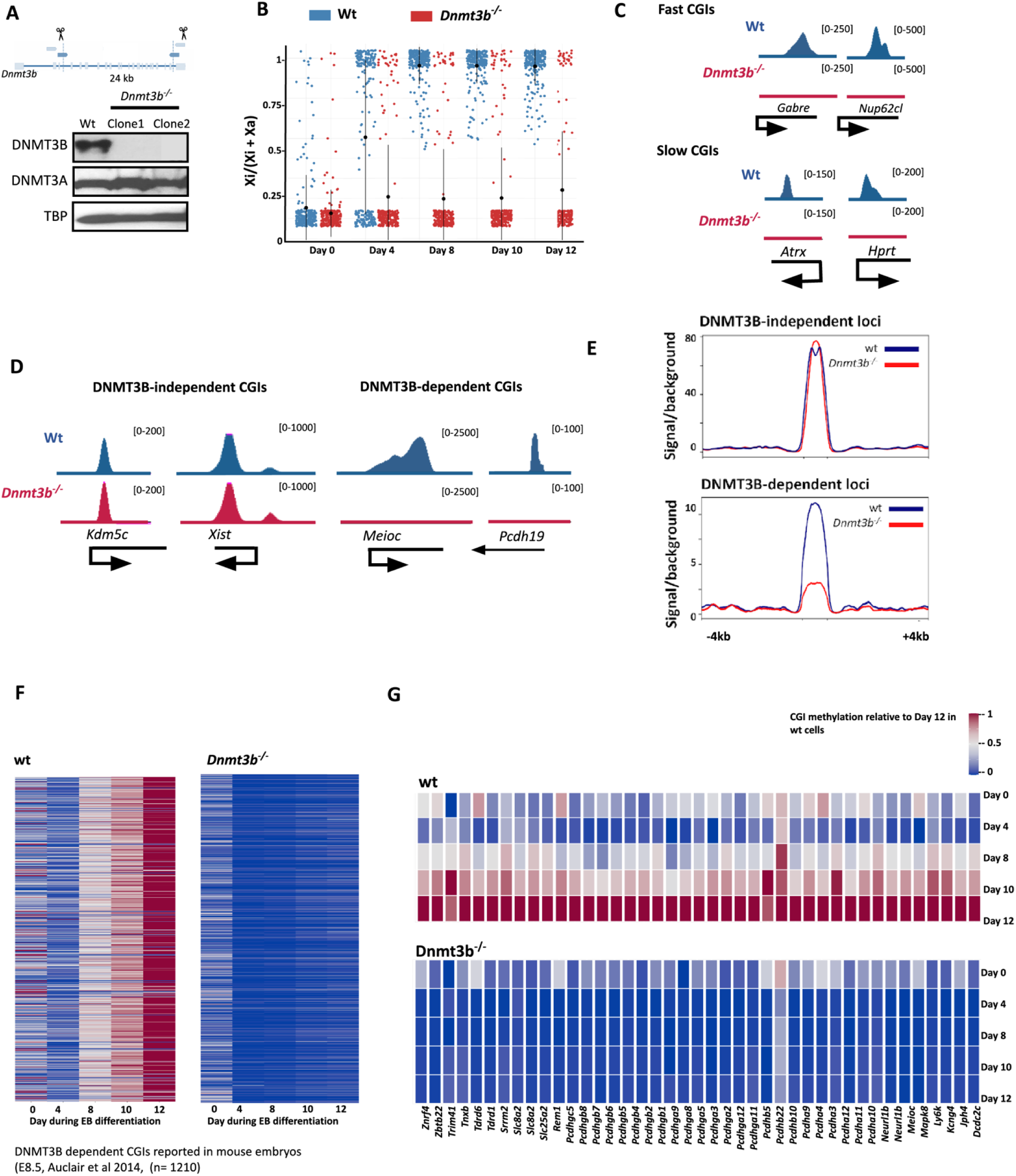
DNMT3B CGI specificity is recapitulated in iXist-ChrX cells. A) Schematic shows the targeting sites of sgRNAs used to generate iXist-ChrX *Dnmt3b^-/-^* cell line (top). Western blot is for two independent Dnmt3b^-/-^ clones. DNMT3A protein levels are not affected in *Dnmt3b^-/-^* cells. TBP loading control. B) Plot showing allelic ratio for all X-linked CGIs in wt and *Dnmt3b^-/-^* iXist-ChrX cells over differentiation timecourse. Most Xi CGIs remain hypomethylated in *Dnmt3b^-/-^* cells. C) Genome browser tracks show examples of MBD-seq data for selected ChrX CGIs. D) Browser tracks illustrating DNMT3B requirement for selected autosomal CGI (*Meioc* and *Pcdh19*) and DNMT3B-independent CGIs. E) Differential peaks analysis for MBD-seq data from wt and *Dnmt3b^-/-^* cells was used to define DNMT3B-dependent and independent loci (for n=5 for wt and *Dnmt3b^-/-^*EBs). Metaplots show MBD-seq data on these peaks in wt and *Dnmt3b^-/-^* EBs. Metaplots show representative data from one of the 5 reproducible replicates for wt and *Dnmt3b^-/-^* EBs (see Supplementary Figure 2). F) Methylation in wt and *Dnmt3b^-/-^* iXist-ChrX CGIs reported to be DNMT3B-dependent in mouse embryo E8.5 (Auclair et al 2014) over a 5-day EB differentiation timecourse. Methylation level relative to EB day 12 in wt iXist-ChrX was plotted for each CGI after spike-in normalization (see methods). G) Heatmap shows methylation levels in wt and *Dnmt3b^-/-^* cells at mouse homologues of CGIs found hypomethylated in *Dnmt3b* mutated ICF patients (Yagi et al,. 2020). As in (F) the methylation levels relative to spike-in normalized EB day 12 in wt iXist-ChrX are plotted.

We also examined genome-wide CGI methylation at developmentally regulated genes previously reported to be DNMT3B-dependent. We performed differential MBD-seq analysis to define DNMT3B-dependent and independent loci (Figure 2D, E and Figure S2G). Levels of CGI methylation increased from d8-d12 of EB differentiation (Figure S2H), a similar trajectory to that seen for slow methylating Xi CGI. Among these CGIs were well-characterized targets, such as germline-expressed genes, protocadherin genes, genes associated with Wnt/β-catenin pathway and *Dppa3* (Auclair et al., 2014; Yagi et al., 2020), all of which were hypomethylated in the differentiated *Dnmt3b^-/-^* iXist-ChrX line (Figure 2F and Figure S2I). Additionally, ICF (immunodeficiency, centromeric instability and facial anomalies), a rare autosomal genetic disorder which predominantly results from mutations in DNMT3B (subtype 1), has been associated with CGI hypomethylation at loci including *Zbtb22, Slc25a31, Slc8a2, Trim42, Ly6k, Tdrd1/6, Rem1, Meioc, Srrm2/4, Znrf4, Jph4, Neurl1b, Mapk8, Kcng4* and *Tnxb* (Yagi et al., 2020). The mouse homologues of these loci were also identified as DNMT3B- dependent CGIs in this analysis (Figure 2G). These results further validate the iXist-ChrX EB differentiation system as a model both for developmentally regulated CGI methylation and Xi CGI methylation.

Interestingly, peak calling on the MBD-seq datasets additionally identified a significant number (6812) of non-CGI annotated peaks. Focusing our analysis on the X chromosome (494/6812 non-CGI peaks), we found that there are two classes of non-CGI MBD elements; one class, that is present both in ES cells and EBs and the other, that gains CpG methylation during differentiation. Among the latter, a subset (n =101) showed gain of methylation specifically on Xi (Figure S3A,B). Non-CGI MBD peaks generally lie >2kb from the nearest TSS and in these instances their methylation is DNMT3B independent. (Figure S3C-E). This contrasts with DNMT3B dependence shown for promoter proximal and CGI peaks (Figure S3C-F). Non-CGI MBD peaks overlapped with regions with high chromatin accessibility (assessed by ATAC-seq) and H3K4me1 ChIP-seq in undifferentiated iXist-ChrX cells, and have a relatively low CG content, suggesting that they likely represent cis-regulatory elements (CREs) (Figure S3G-H). During EB differentiation, these non-CGI MBD elements acquire DNA methylation with fast dynamics (Figure S3I), being fully methylated by day 4 of differentiation.

### CGI methylation by DNMT3B requires the catalytically active isoform DNMT3B1

The *Dnmt3b* gene, unlike *Dnmt3a*, codes for several inactive isoforms that are distinct from their catalytically active counterparts, in both structure and function (Figure S4A). These inactive isoforms have been reported to interact with and regulate catalytic DNMTA/B isoforms (Duymich, Charlet, Yang, Jones, & Liang, 2016). A catalytically inactive isoform of DNMT3B, DNMT3B3, enhances catalytic activity of both DNMT3A and DNMT3B (Gordon et al., 2013; Y. Zeng et al., 2020). Recent work indicates that DNMT3B3 heterodimerizes with catalytic isoforms of DNMT3A/B, and interacts with nucleosomes, so as to place the target recognition domain (TRD) over linker DNA for CpG methylation (Xu et al., 2020). Interestingly, during development DNMT3B3 is upregulated and its expression is maintained in somatic cells (Chedin et al., 2002; Gordon et al., 2013), leading to the suggestion that it is an essential accessory factor fulfilling a similar role to DNMT3L in germline and early embryo cells. Accordingly, loss of CGI methylation in previously described *Dnmt3b^-/-^* ES cells could be attributable to loss of catalytically active DNMT3B1 and/or non-catalytic isoforms, including DNMT3B3 (Figure 3A).

**Figure 3.**
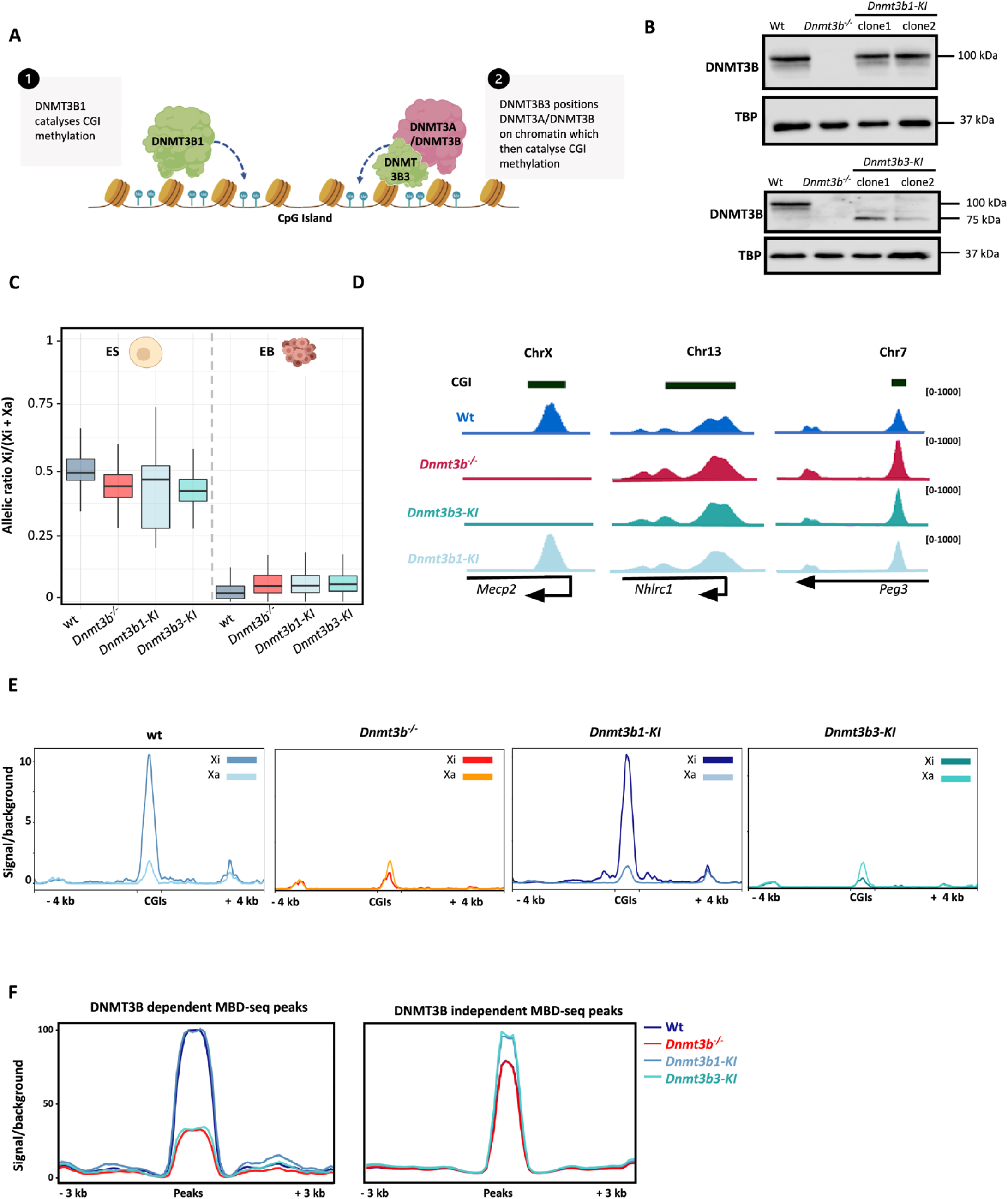
The catalytically active isoform DNMT3B1 is sufficient for developmentally regulated CGI methylation. A) Schematic showing alternative models for DNMT3B-mediated CGI methylation. B) Western blot for two clones for both *Dnmt3b1-KI* (top) and *Dnmt3b3-KI* (bottom). TBP loading control. C) Plot summarising chrRNA-seq results for wt iXist-ChrX, *Dnmt3b^-/-^* and a representative clone for *Dnmt3b1-KI* and *Dnmt3b3-KI* lines on ES cells and EBs. D) Genome browser tracks showing MBD-seq data for selected DNMT3B-dependent CGI (*Mecp2* on ChrX) and two DNMT3B independent autosomal CGIs. E) Metaplots showing MBD-seq signal on Xi and Xa in day 12 EBs in wt, *Dnmt3b^-/-^*, *Dnmt3b1-KI* and *Dnmt3b3-KI* cells. F) Metaplots showing average MBD-seq signal on autosomal DNMT3B-dependent (left) and DNMT3B-independent (right) CGIs in day 12 EBs.

Since DNMT3B isoforms share most of the coding sequence of the gene, engineering mutations of the *Dnmt3b* locus, such that only one isoform is deleted, is not possible (Figure S4A). To overcome this complexity, we used a complementation strategy, in which transgenes encoding specific DNMT3B isoforms were expressed in the *Dnmt3b^-/-^* iXist-ChrX parent line (Figure S4B). For the generation of complementation lines with equivalent expression levels, cDNA encoding DNMT3B isoforms (3B1 or 3B3) were introduced into the TIGRE locus on chromosome 9 (H. Zeng et al., 2008), under a constitutive pCAG promoter, henceforth referred to as *Dnmt3b1* and *Dnmt3b3 knock-in (KI)* lines. Owing to the insertion of sequence encoding Tet-transactivator (rtTA) which allows doxycycline-inducible Xist expression on one allele of the TIGRE locus, the complementation lines carry only a single copy of the respective transgene. Western blot analysis illustrating expression of two independent clones for both *Dnmt3b1-* and *Dnmt3b3-KI* is shown in Figure 3B.

Both the *Dnmt3b1-* and *Dnmt3b3-KI* lines behaved normally with respect to cell differentiation (Figure S4C), and Xi silencing was comparable to the wt iXist-ChrX cells (Figure 3C). Data for independent clones was well correlated (Figure S4D). Analysis of CGI methylation by MBD-seq demonstrates that methylation of DNMT3B-dependent CGIs on Xi is fully rescued in the DNMT3B1 complementation lines (Figure 3D,E). In contrast, there was no detectable Xi CGI methylation on expression of DNMT3B3 (Figure 3D,E). Analysis of CGI methylation genome-wide similarly showed complementation in DNMT3B1 but not DNMT3B3 expressing cell lines, with DNMT3B independent MBD peaks being broadly unchanged (Figure 3D,F). Together, these findings demonstrate that DNMT3B3 is dispensable for the establishment of DNMT3B-dependent Xi and autosomal CGI methylation.

### DNMT3B1 PWWP-ADD domains but not the catalytic domain confer specificity for CGI methylation

Having established that it is the catalytically active DNMT3B1 isoform that is required for Xi CGI, we next explored the role of individual domains in imparting specificity. The catalytic domain of DNMT3 proteins is the most well-conserved domain with 85% of protein sequence conserved. Nevertheless, several studies have shown that subtle differences in the amino acid sequence of DNMT3A and DNMT3B in this region give rise to unique sequence specificities (Emperle et al., 2021; Gao et al., 2020; Handa & Jeltsch, 2005; Lin, Han, Taghva, O’Brien, & Hsieh, 2002; Mallona et al., 2021). To investigate if these differences could underlie DNMT3B1 specificity for methylating CGIs, we complemented *Dnmt3b^-/-^* cells with a chimeric DNMT3B where the catalytic domain of DNMT3B was replaced by that of DNMT3A (represented as *Dnmt3b-3aCat KI*) and performed MBD-seq on 12-day differentiated EBs (Figure 4A). This chimeric construct was able to fully rescue CGI methylation on Xi (Figure 4B- D) and also at DNMT3B-dependent autosomal CGIs (Figure 4E and F). These results demonstrate conclusively that the catalytic domain of DNMT3B does not impart specificity for CGI methylation.

**Figure 4.**
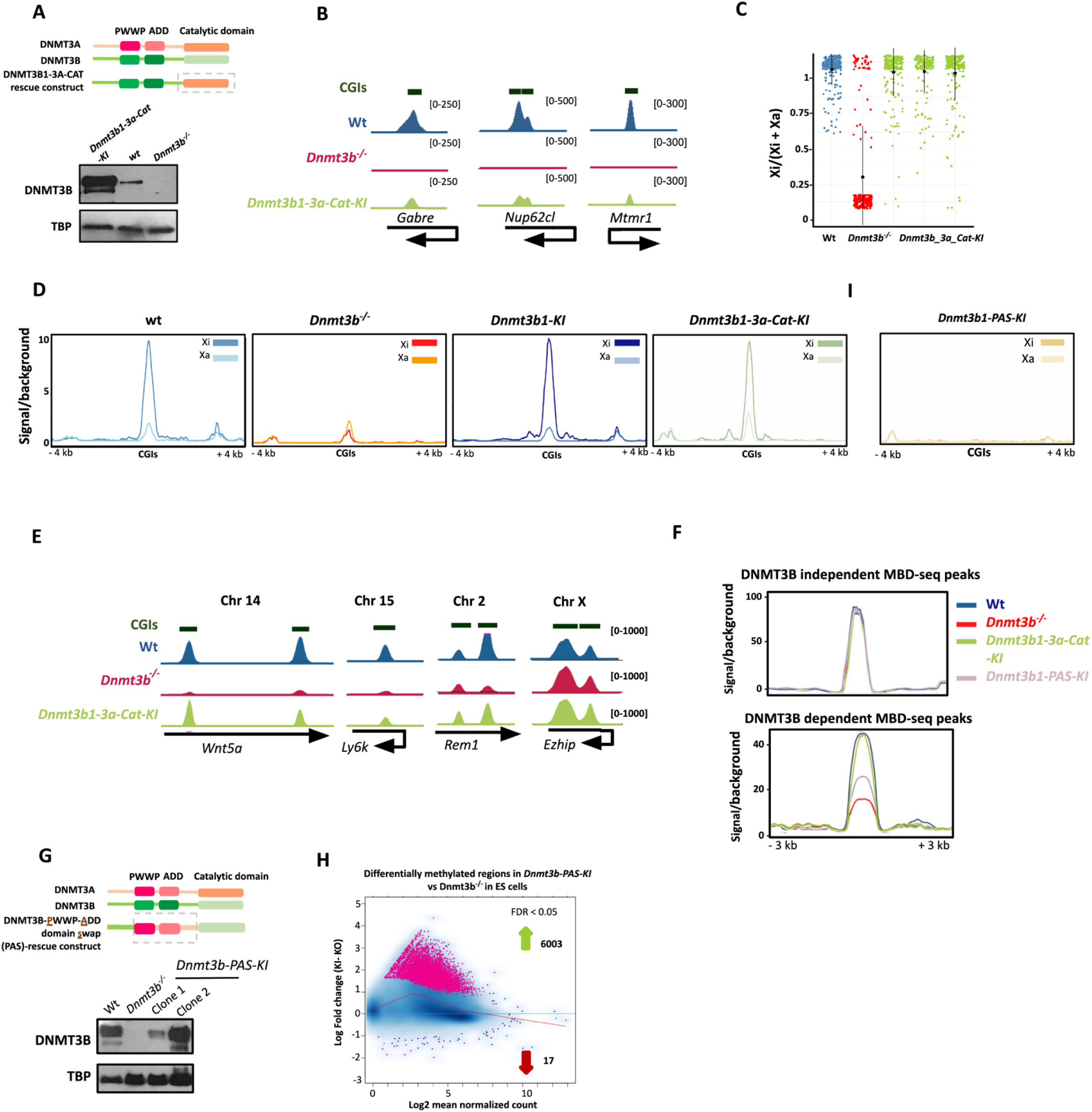
The region of DNMT3B spanning the PWWP and ADD domains imparts specificity for CGI methylation. A) Schematic and western blot illustrating complementation of *Dnmt3b^-/-^* iXist-ChrX cells with a construct encoding a chimeric protein swapping the DNMT3B catalytic domain with that of DNMT3A (*Dnmt3b1-3a-Cat-KI*) B) Genome browser tracks of selected CGIs illustrating DNMT3B1-3A-CAT rescues Xi CGI methylation. C) Plot of allelic ratio for X-linked CGIs in day 12 EBs in wt, *Dnmt3b^-/-^*, three replicates of *Dnmt3b_3aCat-KI* cells. Each dot represents a ChrX CGI. D) Metaplots showing the average MBD-seq signal on Xi and Xa in day 12 EBs in wt, *Dnmt3b^-/-^*, *Dnmt3b1-KI* and *Dnmt3b_3aCat-KI* cells. E) Genome browser tracks showing selected examples of autosomal DNMT3B-dependent CGIs (*Wnt5a* and *Ly6k*) and DNMT3B-independent CGIs (*Ezhip* and *Rem1*). F) Metaplots showing average MBD-seq signal on autosomal DNMT3B-dependent and - independent CGIs in day 12 EBs. DNMT3B1-3A-CAT rescues CGI methylation genome-wide. DNMT3B_PAS rescues a subset of DNMT3B-dependent CGIs. B-F show a representative dataset from 5 reproducible replicates for wt, *Dnmt3b^-/-^*, three replicates of *Dnmt3b_3aCat- KI* cells. G) Schematic and western blot illustrating chimeric DNMT3B protein with region spanning PWWP-ADD domain of DNMT3B swapped with that of DNMT3A and expressed in *Dnmt3b^-/-^* iXist-ChrX cells. H) Plot illustrating that *Dnmt3b_PAS-KI* cells gain methylation (MBD-seq peaks) in ES cells.

The PWWP and ADD domains of DNMT3A and 3B, although largely conserved, have been shown to impart distinct characteristics to the DNMT proteins. The PWWP domain of DNMT3B, for instance, is reported to interact with H3K36me3-modified gene bodies while that of DNMT3A shows preference for H3K36me2-modified intergenic regions (Baubec et al., 2015; Weinberg et al., 2019). Similarly, the ADD domain of DNMT3B has been suggested to form weaker interaction with its catalytic domain compared to DNMT3A and to possess an extended conformation that can serve as an interface for protein-protein interactions (Boyko et al., 2022). Such protein interactors could potentially mediate DNMT3B recruitment to its target loci. To test if the PWWP-ADD region of DNMT3B1 has a role in imparting specificity for CGI methylation we expressed a chimeric DNMT3B with the region spanning PWWP-ADD domains replaced by the equivalent domains from DNMT3A (represented as *Dnmt3b-PWWP- ADD_Swap or Dnmt3b-PAS-KI* cells and performed MBD-seq on 12-day differentiated EBs (Figure 4G and Figure S5A-B). In ES cells, this chimeric DNMT3B protein showed ectopic CpG methylation in comparison to wt iXist-ChrX cells), and ChromHMM analysis revealed that the MBD-seq signal emerged largely from within exons and intergenic regions, indicating that the chimeric DNMT3B protein is catalytically active (Figure 4H and S5C-D). However, most CGIs remained unmethylated both on Xi and autosomes (Figure 4F).

Although most CGIs remained hypomethylated using the PWWP-ADD swap construct, there were some exceptions, totalling around 125 of those CGIs that were DNMT3B- dependent in comparative analysis of wt and *Dnmt3b^-/-^* cells (Figure S5E). Examples of methylated CGIs include *Znf92, Nlgn3* and *Mageb2* on Xi and *Zbtb22* and *Znfr4* reported to be hypomethylated in ICF (Figure S5F). The basis for the differential response of this small subset of CGIs is currently unknown.

### The unique N-terminal domain of DNMT3B is essential for CGI methylation

The N-terminal domains of both DNMT3A and DNMT3B are predicted to be largely flexible unstructured regions and show little if any sequence conservation relative to one another (Figure S6A and B). Despite the lack of conservation between the paralogues (DNMT3A and DNMT3B), the N-terminal domains of both proteins are well conserved among orthologues in primates (Molaro, Malik, & Bourc’his, 2020) (Figure S6C). Several studies indicate that the N-terminal domain is functionally important, but its precise role is not well understood. There is evidence suggesting a function in chromatin binding (Jeong et al., 2009), with involvement of the Polycomb-mediated histone modification H2AK119ub1 in the case of the DNMT3A N-terminal region (Gu et al., 2022; Weinberg et al., 2021), and with the heterochromatin specific protein HP1 in the case of DNMT3B (Taglini et al., 2024) (Figure 5A).

**Figure 5.**
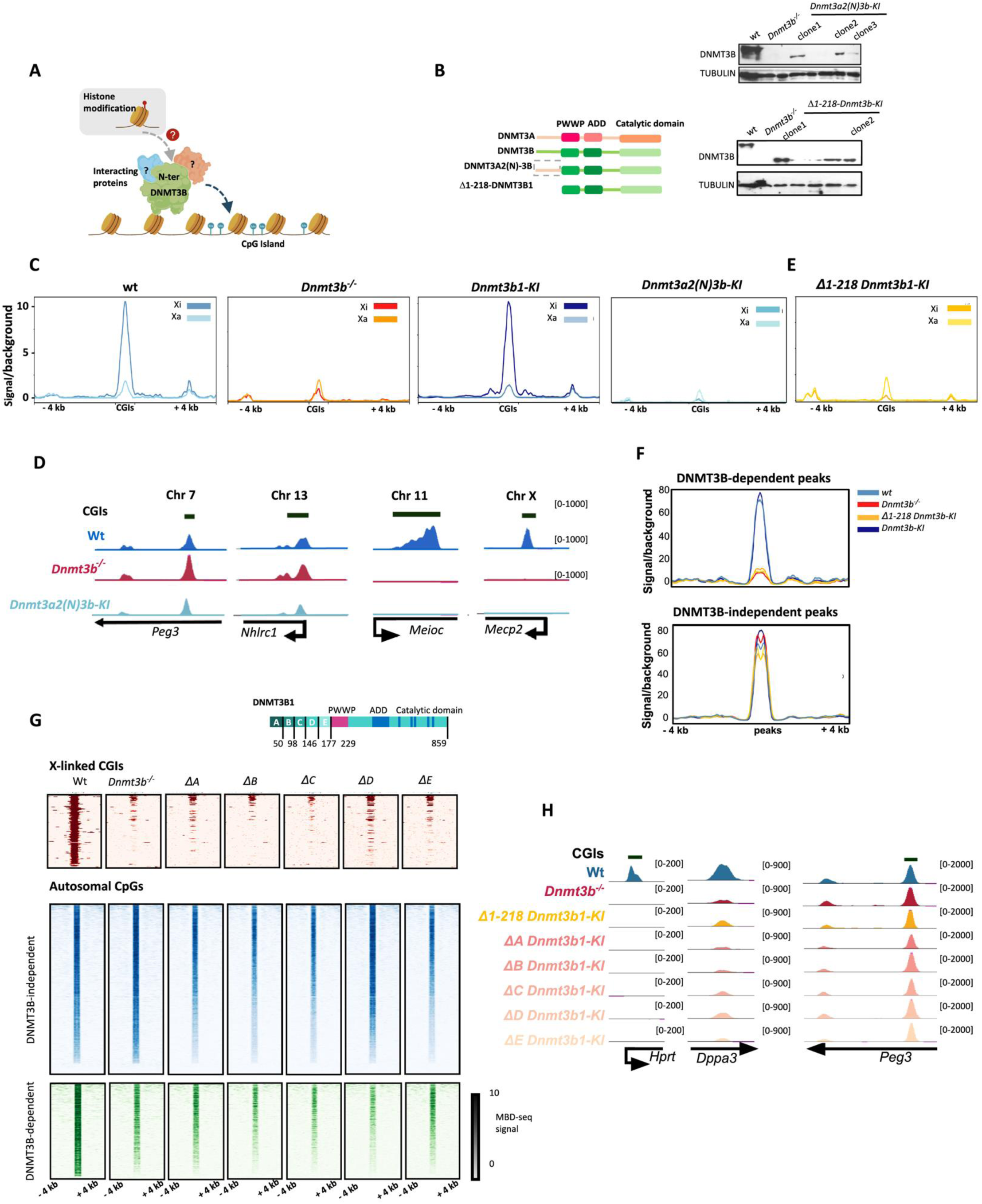
DNMT3B1 N-terminus is required for CGI methylation in *Dnmt3b^-/-^* cells. A) Schematic illustrating hypothesis for a role of the DNMT3b N-terminus in imparting CGI specificity B) Schematic and western blot showing expression of chimeric DNMT3B protein with the N-terminal domain swapped with that from DNMT3A2 (top) or truncation of the N-terminus. C) MBD-seq metaplots showing profiles for methylation of CGIs on the Xa and Xi chromosome following swapping of the N-terminal domain of DNMT3B with that of DNMT3A. *Wt* and *Dnmt3b1-KI* are shown for comparison. D) Genome browser tracks showing selected examples of DNMT3B-independent and DNMT3B- dependent CGIs. E) MBD-seq metaplot showing Xa and Xi CGI methylation for the N-terminal domain truncation mutant. F) MBD-seq metaplots showing methylation of autosomal DNMT3B-dependent (top) and - independent (bottom) CGIs in day 12 EBs in wt, *Dnmt3b^-/-^, Δ1-218 Dnmt3b1-KI* and *Dnmt3b1- KI* cells. G) Schematic (top) of DNMT3B1, highlighting the fragments of the N-terminal domain deleted in *Dnmt3b1* rescue constructs (labelled A-E). Heat maps are across DNMT3B-dependent and independent CGIs on the X chromosome and autosomes as indicated (bottom). Data is from a representative clone of two well correlated complementation cell lines. H) Genome browser tracks showing selected examples of DNMT3B-dependent and DNMT3B-independent CGIs. The complementation lines ΔA-E do not rescue DNMT3B-dependent CGIs.

To investigate if unique features of the N-terminal domain in DNMT3B1 contributes to CGI methylation specificity we expressed a fusion protein replacing the DNMT3B N-terminal region with that from DNMT3A in the Dnmt3^-/-^ iXist-ChrX model (Figure 5B). MBD-seq analysis following EB differentiation revealed complete absence of DNMT3B-dependent Xi and autosomal CGI methylation (Figure 5C-D). To confirm the importance of the N-terminal region, we determined the effect of truncating the DNMT3B1 N-terminus (Δ1-218 DNMT3B1) (Figure 5B). The *Δ1-218 Dnmt3b1-KI* line also failed to rescue Xi and autosomal CGI methylation (Figure 5E,F), whilst DNMT3B independent peaks were retained (Figure 5F).

We went on to establish a series of small N-terminal deletions to determine if there is a specific subdomain of the DNMT3B1 N-terminal region required for CGI methylation (Figure 5G). None of the tested deletions was able to rescue CGI methylation on Xi or autosomes (Figure 5G,H). Deletion of residues 98-146 aa resulted in apparent protein instability (Figure S6D) and accordingly we cannot reach firm conclusions regarding the functional requirement for this sub-region. In sum these experiments demonstrate an absolute requirement for the N-terminal domain of DNMT3B1 for CGI methylation, and moreover indicate that regions within the domain act collaboratively to achieve functionality.

## Discussion

In this study we have established a novel ES cell-based model system for analysing DNA methylation of CGIs on Xi and genome-wide. Key advantages afforded by our model include the use of an inducible promoter to initiate Xist expression, providing for homogeneous and highly synchronised onset and progression of XCI, ease of scale up for analyses requiring larger amounts of material, and amenability to genome engineering to test specified perturbations of key factors. Our system recapitulates key features of CGI methylation using different differentiation procedures, reproducing prior observations such as distinct early and late Xi CGI methylation phases during cell differentiation, differentiation-dependent methylation of a defined specific subset of autosomal CGIs, and DNMT3B dependence of CGI methylation on Xi and autosomes. In addition to confirming prior findings, our initial MBD-seq analysis revealed that a subset of non-CGI elements that correlate with distant CREs gain CpG methylation on Xi in a DNMT3B-independent manner. This observation may help to explain a prior observation that loss of DNMT3B1 alone does not result in significant derepression of Xi genes during embryogenesis, contrasting with treatment of somatic cells with demethylating agents which can result in sporadic Xi gene reactivation. Further studies are required to determine the precise contribution of CRE CpG methylation on Xi in establishment and/or maintenance of X chromosome inactivation.

Several studies have highlighted that DNMT3 proteins form homo- or heterodimers (Xu et al., 2020; Y. Zeng et al., 2020), with heterodimers involving non-catalytic DNMT3L being important in germ cells and undifferentiated embryonic cells (Bourc’his, Xu, Lin, Bollman, & Bestor, 2001; Ooi et al., 2007; Suetake et al., 2004), and DNMT3B3 implicated in somatic lineages (Xu et al., 2020; Y. Zeng et al., 2020). We used complementation analysis in a *Dnmt3b^-/-^*background to demonstrate that the catalytic DNMT3B1 isoform alone (possibly as heterodimer with DNMT3A) is sufficient for CGI methylation on the Xi and autosomes and that DNMT3B3 alone, despite being available to heterodimerise with DNMT3A, could not restore this activity. These observations pinpoint CGI methylation specificity to the catalytically active DNMT3B1 isoform, both in XCI and genome-wide. We note that MBD-seq analysis did not detect CpG methylation changes reported previously using limited representation bisulfite analysis of mouse embryos expressing a non-catalytic DNMT3B mutant (Nowialis et al., 2019). It may be that these differences require higher resolution than that provided by MBD-seq and in this regard it would be interesting to examine global CpG methylation at single nucleotide resolution in our ES cell model using newly developed long-read sequencing methods.

Given our finding that the DNMT3B1 isoform alone is sufficient for CGI methylation, we were able to set about defining which parts of the protein are required for this activity. Our DNMT3A domain swap experiments ruled out unique features of the DNMT3B1 catalytic domain, instead implicating the PWWP-ADD and N-terminal domains as being important for CGI specificity. In addition to the N-terminal truncation mutant, expression of a series of smaller N-terminal deletions also failed to restore CGI methylation, further verifying this finding. We speculate that the DNMT3B1 PWWP-ADD and N-terminal regions synergise in recognising an interacting protein or a specific chromatin feature/post-translational modification, that underlies CGI specificity. An intriguing possibility relates to the histone PTM H3K9me3 which is enhanced/redistributed on Xi following recruitment of the chromosomal protein SmcHD1(Ichihara, Nagao, Sakaguchi, Obuse, & Sado, 2022). SmcHD1 in turn is required for CpG methylation at the majority (∼90%) of Xi CGIs (Blewitt et al., 2008; Gendrel et al., 2012). Potential mechanisms include direct binding of DNMT3B to H3K9me3, and here it is interesting to note that the protein ATRX has an ADD domain which has been shown to mediate H3K9me3 binding (Iwase et al., 2011). Further studies are required to investigate this and alternative hypotheses.

## Methods

### Cell culture conditions

Mouse ES cells (iXist-ChrX cells (Nesterova et al., 2019), and the derived cell lines) were cultured in Dulbecco’s Modified Eagle Medium (DMEM, Gibco) supplemented with 10% foetal calf serum, 2 mM L- glutamine, 0.1 mM non-essential amino acids, 50 µM 2-mercaptoethanol, 100 U/ml penicillin, 100 µg/ml streptomycin (all from Gibco) and leukaemia inhibitory factor (LIF, made in-house), under standard culture conditions (37°C, 5% CO2, humid). ES cells were seeded atop a monolayer culture of inactivated mouse SNL fibroblast cells (previously treated with Mitomycin C, referred to as “feeder cells”), on gelatinised tissue culture plates. Gelatine coating of plates was achieved by treatment with 0.1% gelatine in PBS (Gibco) for 2- 3 minutes at room temperature. Upon attaining 80% confluency, cells were detached using TrypLE Express (Gibco) and passaged. While in culture, cells were fed with LIF supplemented ES medium every day.

### Generation of cell lines

sgRNA for the CRISPR/Cas9 mediated double stranded breaks at the target loci were for generating *Dnmt3b^-/-^* cells and for targeting of DNMT3B rescue constructs in TIGRE locus were designed using online tool CRISPOR. sgRNAs were cloned into pX459 plasmid (Addgene plasmid #62988) using a single step digestion-ligation protocol developed by the Zhang lab (Ran et al., 2013). For the generation of complementation cell lines, a donor plasmid with a constitutive CAG promoter, 1000 bp long homology arms for TIGRE locus and puromycin resistance to allow downstream selection of successfully targeted cells was used (gifted to the lab (Nora et al., 2017)). Full length DNMT3B or truncated variants of DNMT3B were PCR amplified from cDNA using a high fidelity Phusion DNA polymerase (Invitrogen) as per manufacturer’s guidelines. The PCR amplified DNA was extracted from the agarose gel using Zymoclean gel recovery kit (Zymo Research) and cloned into the TIGRE targeting plasmid using Gibson assembly to create a ‘donor homology construct’ for generation of stable cell lines. Gibson assembly was performed using NEB master mix, following the protocol recommended by the manufacturer. The plasmids were transformed into XL10-Gold Ultracompetent cells using protocol recommended by Stratagene. After cloning, the sequence of the Dnmt3b donor homology construct plasmid was verified using Sanger sequencing.

Prior to transfection, cells were plated in 6 well plates with feeders for 24 hours. An hour before transfection, the medium was replaced by an antibiotic-free medium (i.e., without penicillin and streptomycin). Lipofectamine 2000 (Life Technologies, Invitrogen) was used for transfection. For generation of the knockout lines, 1 µg each of the pX459 plasmids with CRISPR guides cloned for targeting the 5’ and 3’ cleavage sites were transfected. For the knock-in lines the “donor homology construct” plasmid and pX459 plasmid with cloned sgRNA for targeting TIGRE were co-transfected in a molar ration of 6:1 (3.5 µg of the plasmid with the donor homology construct and 1 µg of the pX459 with cloned guide RNA). Lipofectamine complexes were allowed to form in serum-free Opti-MEM (Gibco), as per manufacturer’s instructions. 24 hours after transfection, the cells were trypsinized and split into 90 mm gelatinized dishes with feeders in ratio 1/3, 1/10 and 1/15, and fed with 2.5 µg/ml puromycin-supplemented medium to allow for selection of transfected cells. Cells were maintained in the selection medium for 2 days and then later were switched to regular cell culture medium, till the colonies appeared (around 10 days later). Single colonies were picked in 96-well plates and split 1/3 upon reaching 80% confluency to give 3 replica plates which were used for expansion, screening and freezing of the clones.

### Embryoid body (EB) differentiation

ES cells were pre-plated to separate and remove the feeder cells from ES cells in EC10 medium. EC10 medium had the same composition as the medium used for culturing ES (described above) but does not contain LIF. After pre-plating, the cell suspension was moved to a falcon tube and the cells were counted using LUNA-II automated counter (Logos Biosystems). For setting up EB differentiation, 1 x 10^6^ cells were plated in 10 cm, ungelatinized petri dish containing 10 ml of EC10 medium supplemented with 1.5 µg/ml doxycycline. For the next two days (day 1 and day 2), cells were fed with EC10 containing 1.5 µg/ml doxycycline; at this stage cells made small sparse colonies. On day 3, the small colonies were gently blown using a pipette to dislodge the weakly attached colonies from the plate. The suspended colonies were then moved to 10 cm bacterial culture dishes, which owing to their different material, provides a less sticky surface. Until day 7, these colonies stay suspended and form spherical masses called embryoid bodies (EBs). From day 3 to day 7 EBs were fed with EC10 + 1.5 µg/ml doxycycline by taking the contents of the plate (medium with suspended EBs) out in a 50 ml falcon tube, allowing the EBs to settle (usually within 5 minutes, the exact time depends on the size of EBs; when settled EBs form a loose pellet), aspirating the old medium, gently resuspending EBs in fresh medium, and placing them back in the culture dish. On day 7, the EBs start to attach to the dish. The weakly attached EBs were carefully detached by pipetting and moved back to 10 cm cell culture dishes. From day 7 to day 11 these attached EBs were fed every day. On day 12, the EBs were collected, counted, pelleted, and used for further experiments. For certain experiments, cells were collected at timepoints within this full timecourse such as day 4, day 8 and day 10.

### Methyl binding domain affinity purification (MBD)

MBD-affinity purification was performed using the Epimark Methylated DNA Enrichment kit manufactured by NEB (Cat no. E2600S). The protocol was carried out as per the manufacturer’s guidelines. Genomic DNA was sonicated to a fragment size range of 100-300 bp at 4°C in TE buffer using Diagenode Bioruptor Pico. 5 µg of sonicated genomic DNA was used as input and mixed with 150 ng of DNA from HeLa cells (3% of input DNA from mouse cells) which served as an internal calibration for downstream quantitative bioinformatic analysis. Genomic DNA from Hela cells was isolated and sonicated in bulk and the same stock was used as control for all MBD experiments. Enrichment of methylated DNA was then validated using qPCR analysis using primers targeted against CpG islands of imprinted loci Daz1 and Ezhip which are methylated in our model cell lines (primer sequences are given). The qPCR was performed using a 20 µl reaction containing 2X SensiFAST SYBR No-ROX master mix (Bioline), 10 µM of primers and diluted DNA (1:50 dilution in nuclease free water) from input and immunoprecipitated (IP) fraction using Rotor-Gene Q machines (Qiagen). The qPCR program details and the primers used for qPCR validation are given below:

Daz1_CGI_F: GCGTTATCCCGTCAGCAAAG

Daz1_CGI_R: CAAGGCCTGCTCTCTAGTCGT

Ezhip_CGI_1_F: AGCGGAGGGCCTCGG

Ezhip_CGI_1_R: CCTGACCAAAGCCACCAT

Ezhip_CGI_2_F: TGCTCCTACGACCGAAAGG

Ezhip_CGI_2_R: CCCGATGATGATGATGACG

| Cycles | Temperature (°C) | Time | Step |
| --- | --- | --- | --- |
| 1 | 95 | 2-3 min | Polymerase activation |
| 40 | 95 | 10 s | Denaturation |
|  | 60 | 10 s | Annealing |
|  | 72 | 20 s | Extension |
| Hold | 50 | 10 min | - |

### NGS library preparation, validation, and quantification

The fragment size for the pulled down DNA after MBD, BioCAP and ChIP experiments was checked using Bioanalyzer 2100 (Agilent) with High sensitivity DNA Chip protocol to ensure that the fragments were within the range of 150-800 bp. If larger, the DNA was sonicated using Diagenode Bioruptor Pico (detailed above). DNA was quantified using Qubit double stranded DNA high sensitivity kit (Thermofisher) for batch library preparation. Quantity of starting/input DNA from the samples was diluted in TE buffer to be within the same range (± 10 ng). NGS DNA library was prepared using NEBNext Ultra II DNA library preparation kit from Illumina as per manufacturer’s recommendations. After library preparation, the libraries were quantified using Qubit and diluted to 2 ng/µl in TE buffer. Size of the libraries was checked to be between 300 and 1000 bp, by running them on Bioanalyser. Libraries were quantified using qPCR with KAPA library quantification DNA standards (Roche) before pooling (using a 20 µl reaction containing 2X SensiFAST SYBR No-ROX master mix (Bioline), 20 µM of Illumina primers and 1:2000 dilution of 2 ng/µl library on Rotor-Gene Q machines (Qiagen), program and primer sequences are in Table). After pooling, the library pool was again quantified with qPCR alongside the library pools from sequenced NGS runs to serve as a reference. The libraries were sequenced using 150 cycle, paired-end NextSeq 500/550 v2.5 Kits from Illumina at University of Oxford, Department of Zoology Nextseq sequencing facility on Illumina NextSeq500 (FC-404-2002).

### Chromatin RNA extraction and library preparation for sequencing (chrRNA-seq)

Cell pellets (1 x 10^5 – 3 x 10^6) were washed with PBS (Gibco), resuspended in RLB (10 mM Tris pH 7.5, 10 mM KCl, 1.5 mM MgCl_2_, and 0.1% NP-40) and incubated on ice for 5 minutes. RLB with 24% sucrose was then added to the suspended pellets and cytoplasmic fraction was separated from the nuclei by centrifugation at 2800 x g for 10 minutes at 4o C. The nuclei were resuspended in NUN1 (20 mM Tris pH 7.5, 75 mM NaCl, 0.5 mM EDTA, 50% glycerol, 0.1 mM DTT). Nuclear lysis was conducted by addition of equal volume of NUN2 (20 mM HEPES pH 7.9, 300 mM, 7.5 mM MgCl_2_, 0.2 mM EDTA, 1 M Urea, 0.1 mM DTT) and incubation for 15 minutes on ice with occasional mixing. The chromatin fraction was isolated as an insoluble pellet by centrifugation at 2800 g for 10 minutes at 4°C. The RNA was extracted by suspending and homogenizing chromatin pellet in TRIzol (Invitrogen). RNA was then extracted using chloroform and precipitated with isopropanol. RNA pellet was washed with 70% ethanol and resuspended in nuclease free water. RNA was then treated with Turbo DNAse (Thermofisher scientific) and quantified using Qubit RNA HS Assay Kit. This RNA was used for NGS library preparation using TruSeq stranded total RNA kit from Illumina as per manufacturer’s recommended protocol.

### Nuclear extract preparation

10-20 x 10^6 cells were pelleted and washed with PBS. The cell pellet was resuspended in 10 volumes of cold buffer A (10 mM HEPES pH 7.9, 1.5 mM MgCl_2_, 10 mM KCl, supplemented with 0.5 mM PMSF, 0.5 mM DTT, and 1X complete EDTA-free proteinase inhibitor (Roche), before use). The resuspended pellets were incubated on ice for 10 minutes and centrifuged at 1500 g for 5 minutes at 4o C. Pellets were again resuspended in 3 volumes of buffer A with 0.1% NP-40 and again incubated on ice for 10 minutes with occasional inversion. Nuclei were recovered with centrifugation (400 g for 5 minutes at 4°C). The pelleted nuclei were then resuspended in 1 volume of buffer C (250 mM NaCl, 5 mM HEPES pH 7.9, 26% glycerol, 1.5 mM MgCl_2_, 0.2 mM EDTA pH 8, with freshly added 0.5 mM DTT and complete EDTA-free PIC). 5 M NaCl was added to the lid of the 1.5 ml tube so that the final concentration of NaCl in the extract (resuspended nuclei and buffer C) becomes 400 mM and mixed by quickly inverting to avoid any precipitation. The extraction was allowed to proceed for 1 hour at 4o C. The nuclear extracts (liquid fraction) were recovered using centrifugation (16000 g for 20 minutes at 4°C). The nuclear extract was quantified using Bradford assay (BioRad) and 25 µg of protein was used for western blot analysis.

### MBD-seq data analysis

#### Mapping of fastq files for MBD-seq

Nextseq fastq files were mounted on the Oxford Biochemistry genomics server using Basemount from Illumina. For calibrated MBD-seq, the fastq files were mapped to N-masked mm10 genome concatenated with human hg38 genome using bowtie2 V2.3.5.1 (Langmead & Salzberg, 2012) with “--sensitive”, “--no-mixed”, “--no-discordant” and “--maxins” set to 800. PCR duplicates were removed using picard-tool “MarkDuplicates” script. The aligned reads were then split among parental genomes*, M.musculus Castaneous* (Cast) or *Mus.musculus Domesticus* (129S) with SNPsplit (Krueger & Andrews, 2016) using “paired” parameter and SNP file containing the information regarding the 23,005,850 SNPs between Cast and 129S genomes (UCSC, provided by Dr. Guifeng Wei in the lab). For visualization of data normalized bigwig files were generated using deeptools “bamCoverage” with the parameter “scaleFactor (1 /read count)” where readcount is the number of reads successfully aligned to the reference genome for each sample, counted using samtools.

#### Counting reads in CGIs, normalization of MBD-seq data using spiked-in human DNA and allelic ratio calculation

FeatureCounts V1.5.2 (Liao, Smyth, & Shi, 2014) was used to count the number of reads overlapping regions of interest, for instance CGIs (annotation used was from publication (Illingworth et al., 2010). the number of reads mapping to human and mouse genome were counted using samtools. A “normalization factor” was calculated using the formula derived and described in detail by (Hu et al., 2015). For clarity, the equation used is given below:

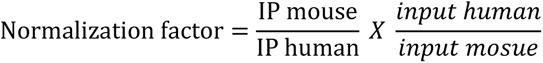

Where:

- IP mouse = number of reads in MBD-/ChIP-seq sample mapped to mouse genome
- IP human = number of reads in MBD-/ChIP-seq sample mapped to human genome
- Input mouse = number of reads in corresponding input sample mapped to mouse genome
- Input human = number of reads in corresponding input sample mapped to human genome

For every comparison, datasets were further normalized to a control. For example, when comparing the relative gain in CGI methylation during EB timecourse, ES cells at day 0 were considered as control with normalization factor 1, and other timepoints (day 4, day 8, day 10 and day 12) were normalized to day 0.

For allelic ratio calculation, the reads overlapping the CGIs were counted via featureCounts on allelically split bamfiles. The CGIs covered in low sequencing depth (i.e. Xi + Xa < 7) were removed. The allelic ratio of filtered CGIs was then calculated (Xi/ (Xi +Xa)). For MBD-seq, unmethylated CGIs on ES cells gave no reads as the assay only captures methylated CpGs (i.e., Xi + Xa = 0). Therefore, the filter of (Xi+ Xa ≥ 7) was applied at the last three time points of differentiation timecourse.

MBD-seq data for different cell lines was collected from at least three replicates, collected either from three individual clones (*Dnmt3a2(N)-3b-KI* and *Dnmt3b-PAS-KI*) or two clones with two technical replicates from one of the clones (*Dnmt3b*^-/-^, *Dnmt3b1-KI, Dnmt3b3-KI*, *Δ1-218 Dnmt3b1-KI* and *ΔA-E Dnmt3b1-KI* lines). Only one clone was obtained for *Dnmt3b-3a(Cat)- KI* line so the MBD-seq data was obtained from three technical replicates. Of the three clones *Dnmt3b-PAS-KI* cell line, two clones lost one X chromosome owing to which the allelic analysis only represents data from one clone.

#### Finding differentially methylated regions between two conditions

Differentially methylated regions in this work were the differential MBD-seq peaks in the two tested conditions (i.e., ES versus EBs or wt vs knockout). This analysis was performed using a bioconductor package called DiffBind (Stark & Brown, 2011). MACS2 V2.2.7.1 (Zhang et al., 2008) was used to call peaks with parameters for paired-end bam file and q value of 0.01 (“- f BAMPE” and “-q 0.01”). DiffBind analysis was performed using the MACS called peaks as recommended by the authors of DiffBind using DESeq2 script inbuilt in DiffBind (Stark & Brown, 2011). The differential regions (MBD-seq peaks in this case) were defined as statistically significant if the false discovery rate (FDR) was less than 0.01. The log2 fold changes and FDR values were used to plot volcano and MA plots.

#### Finding genes associated with CGIs

CGIs (Illingworth et al., 2010), were allotted to the closest gene using a HOMER software (Illingworth et al., 2010) command “annotatePeaks.pl” which provided distance to the closest gene (as defined by RefSeq, (Pruitt, Tatusova, & Maglott, 2005), and the genomic feature (exon, intron, promoter, intergenic etc) for every individual CGI.

#### Generation of heatmaps and profile plots

Profile plots (or metaplots) for the visualization of average MBD- or BioCAP-seq signal at CGIs was generated using deeptools V3.5.1 (Ramírez, Dündar, Diehl, Grüning, & Manke, 2014). The “computeMatrix scale-regions” command with parameter “binSize 50” on size normalized bigWig file was used. The output from computeMatrix was used to run “plotProfile” or “plotHeatmap” using “region file” containing the genomic coordinates for the regions of interest (e.g., CGIs). In this work one profile plots or heatmaps of one representative replicate out of 3 highly correlated replicates are shown. The reproducibility among the replicates is shown by scatter plots in the supplementary figures or PCA plots.

### Mapping chrRNA-seq fastq files

The procedure for mapping chrRNA-seq fastq files has been detailed in (Nesterova et al., 2019). The paired-end reads aligning to mm10 rRNA sequence using bowtie2 (Langmead & Salzberg, 2012) were discarded. The remaining reads were then mapped to N-masked mm10 genome using STAR (Dobin et al., 2013) with “--outFilterMultimapNmax 1”, “-- outFilterMismatchNoverReadLmax 0.06”, “ --alignEndsType EndToEnd”. As for MBD-seq d, the aligned reads were assigned to *M.musculus Castaneous* (Cast) or *Mus.musculus Domesticus* (129S) genomes using SNPsplit (Krueger & Andrews, 2016). Reads overlapping genes (Refseq annotation (Pruitt et al., 2005)), were counted and assigned to the given genes using featureCounts (Liao et al., 2014). DESeq2 was used to assess the statistically significant changes in the gene expression between various cell lines (for example wt and *Dnmt3b^-/-^* lines) (Love, Huber, & Anders, 2014). The parameter p-adj < 0.05 was used to define the threshold for a gene to be called significantly differentially expressed. The results for this analysis were visualized using MA and volcano plots generated using R and ggplot2.

## Acknowledgements

We would like to thank members of the lab for discussion and suggestions for this project, and Dr Heather Coker for critical reading of the manuscript. All work was funded by Wellcome Trust (Grant number 215513) and a Wellcome Trust PhD studentship awarded to Komal Yasmin (Grant number 215109).

---

**Supplementary Figure 1.**
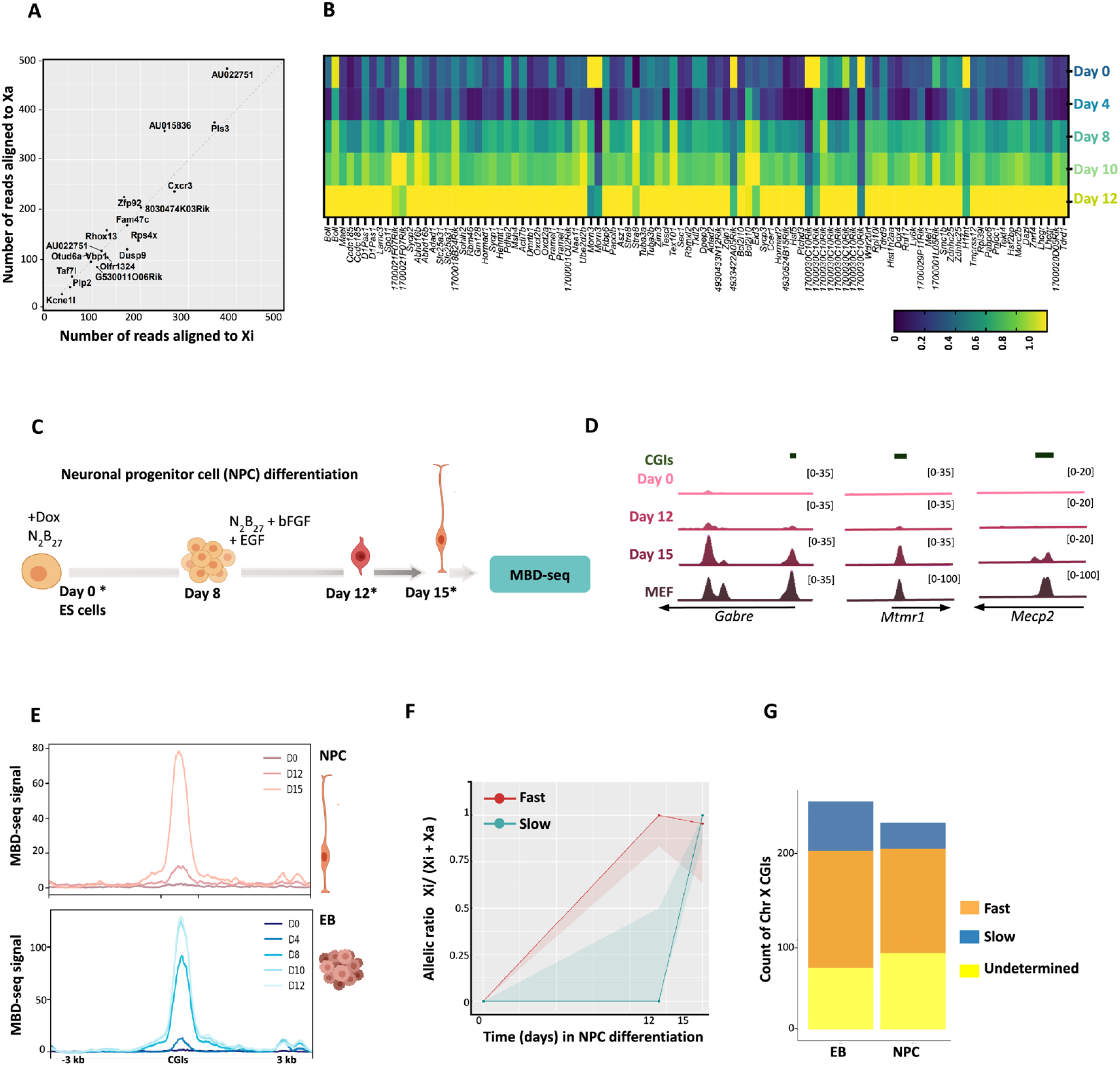
EB differentiation of iXist-ChrX cells models developmentally regulated CGI methylation. A) Dot plot showing CGIs that have MBD-seq signal on both Xi and Xa. Germline-specific genes such as Rhox13, Taf7l, AU022751 fall into this category. B) Heatmap shows the gain of methylation (measured by MBD-seq) on germline genes (Auclair et al., 2014; Endoh et al., 2017) during the 5-day EB timecourse. The methylation level relative to spike-in normalized EB day 12 is shown for each CGI. C) Schematic illustrating NPC differentiation method. Asterisks indicate the days at which cells were collected for the experiment. D) Tracks show MBD-seq signal on chrX CGIs at three time points during NPC differentiation. MBD- seq data for mouse embryonic fibroblasts (MEF) in which X inactivation is established is shown for reference. E) MBD-seq metaplots showing the average profile of CGIs during NPC and EB differentiation. D0- D15 represents day 0 -day 15 of differentiation. F) Line plots showing the allelic ratio for “fast” (red) and “slow” (green) CGIs during NPC differentiation. The solid line shows the median while the shaded ribbon shows the interquartile range for allelic ratio. X-axis shows time in days during differentiation. G) Counts of CGIs assigned to “fast” and “slow” methylating categories in NPC and EB differentiation systems.

**Supplementary Figure 2.**
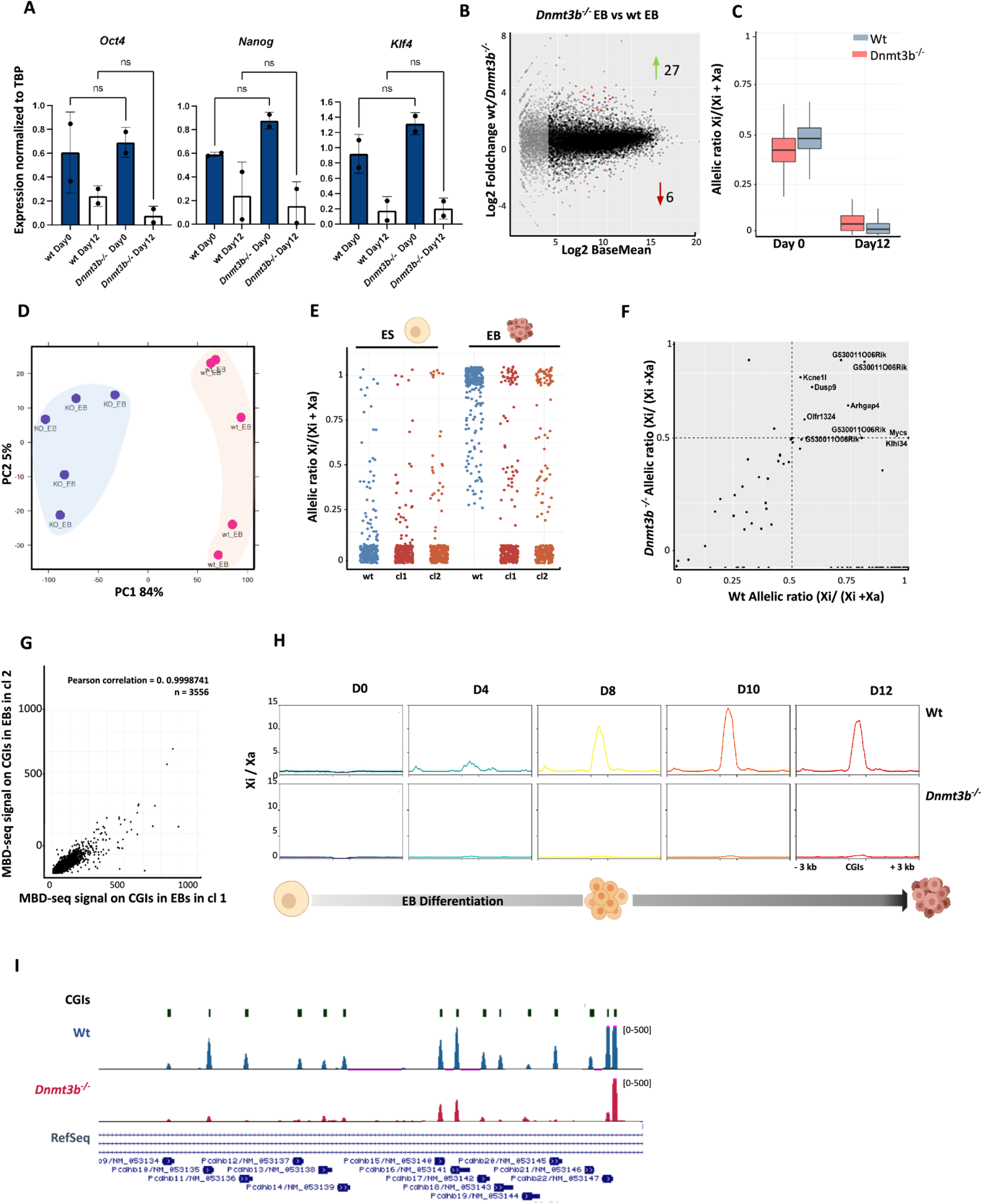
DNMT3B mediates developmentally regulated CGI methylation. A) Plots showing the expression levels of pluripotency genes *Oct4, Nanog*, and *Klf4* (downregulated through differentiation) in both wt and *Dnmt3b^-/-^* (KO). Expression levels were normalized to the housekeeping gene TATA-box binding protein (TBP). Graphs summarize data from two technical replicates for *Dnmt3b^-/-^*and wt cells. B) MA plot comparing gene expression between *Dnmt3b^-/-^*and wt EBs. The significantly up- and down-regulated genes are shown as red dots and the counts are shown beside the arrows. C) Boxplots showing allelic ratio for chromatin RNA-seq data in wt and *Dnmt3b^-/-^* cells before and after differentiation (day 0 and day 12). D) PCA plot shows the reproducibility among MBD-seq replicates for wt and *Dnmt3b^-/-^* cell lines differentiated to EBs. E) Plots showing the allelic ratio at X-linked CGIs for wt and two clones of *Dnmt3b^-/-^* cells, before and after 12-day EB differentiation (ES and EB respectively). Wt is representative example from 5 replicates. F) Allelic ratio of X-linked CGIs in wt plotted against *Dnmt3b^-/-^*line in day 12 EBs, each dot equates to one CGI. CGIs on the X chromosome that gain methylation independent of DNMT3B are labelled. Gain of methylation specifically on Xi is indicated by allelic ratio > 0.5 (upper right quadrant). G) Dot plot showing correlation between MBD-seq data obtained from 12-day differentiation of two different *Dnmt3b^-/-^*clones. H) MBD-seq metaplots illustrating gain of CGI methylation on the Xi at indicated differentiation timepoints in wt and *Dnmt3b^-/-^*lines. D0-D12 represent day 0 – day 12 of EB differentiation. I) Genome browser view of MBD-seq signal in wt and *Dnmt3b^-/-^* EBs at *Pcdh* cluster CGIs which are methylated in a DNMT3B dependent manner.

**Supplementary Figure 3.**
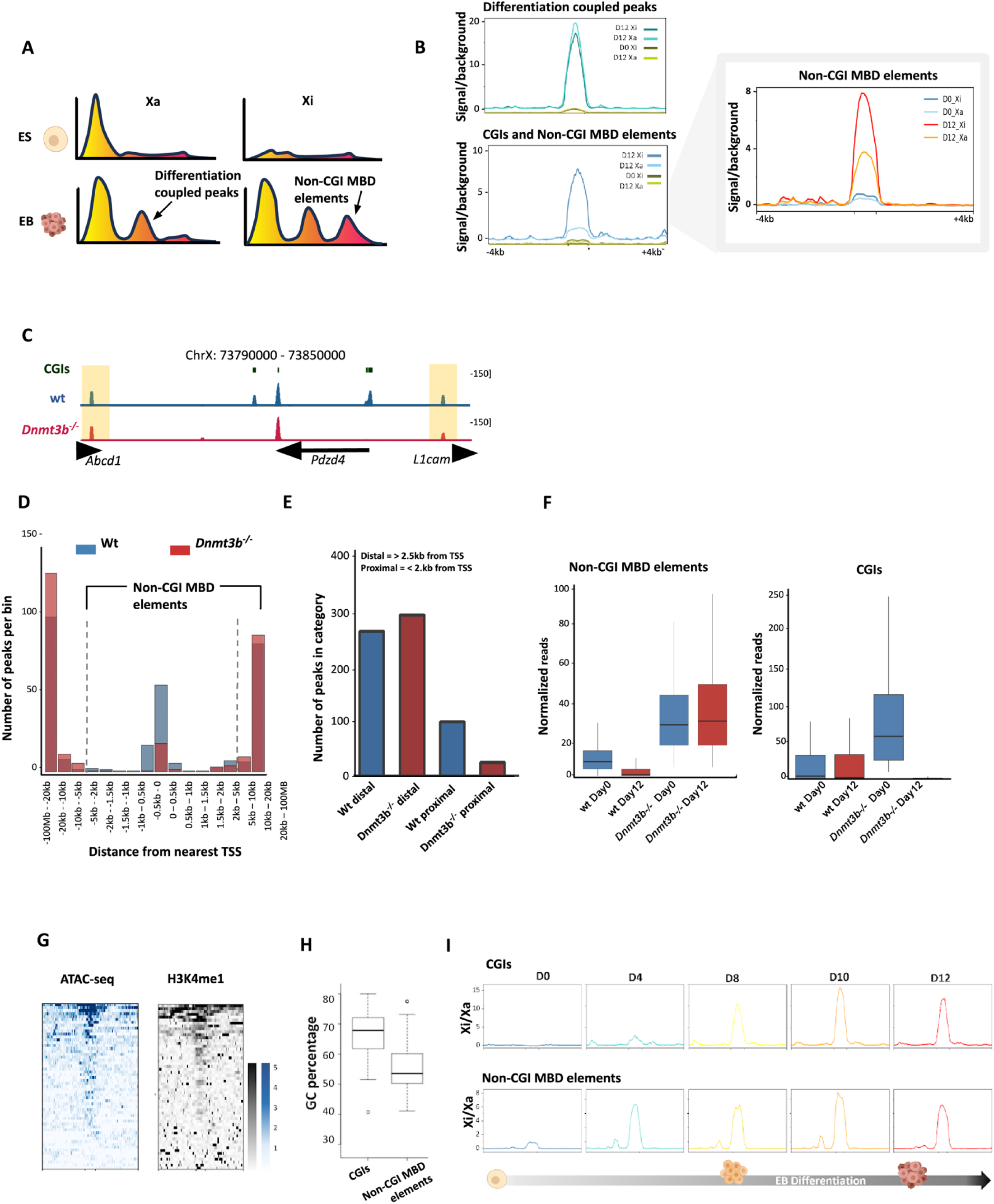
Non-CGI MBD elements represent putative CREs that gain methylation on Xi independent of DNMT3B. A) Schematic showing the strategy for categorizing MBD-seq peaks into differentiation-coupled and non-CGI MBD elements. The later specifically gain methylation on Xi. B) MBD-seq metaplots for Xi and Xa at two classes of loci defined in (A). Inset is a metaplot that shows the average MBD-seq signal on Xi and Xa at non-CGI MBD elements at day 0 (ES) and day 12 EBs. non-CGI MBD elements show allelic bias with MBD-seq signal largely originating from Xi. C) Genome browser tracks showing the novel “non-CGI MBD elements” highlighted in yellow. Non-CGI MBD elements reside distant from gene promoters (>2 kb) and gain DNA methylation independent of DNMT3B during EB differentiation. D) Plot shows the counts of MBD-seq peaks at chrX at various distance from TSS. *Dnmt3b* deletion mainly impacts CGIs (which lie within 2kb of TSS). E) Counts of MBD-seq peaks (categorized into promoter distal or proximal) in wt and *Dnmt3b^-/ -^*EBs. F) Plot comparing the methylation levels (expressed as normalized reads from calibrated MBD- seq) at CGIs and non-CGI MBD elements in wt and *Dnmt3b^-/-^* EBs. G) Heat maps illustrating that non-CGI MBD elements correlate with chromatin regions of high accessibility and H3K4me1 deposition. H) CG percentage at non-CGI MBD elements. I) Non-CGIs elements exhibit fast methylation dynamic with large allelic bias (Xi/Xa signal) at day 4 on average.

**Supplementary Figure 4.**
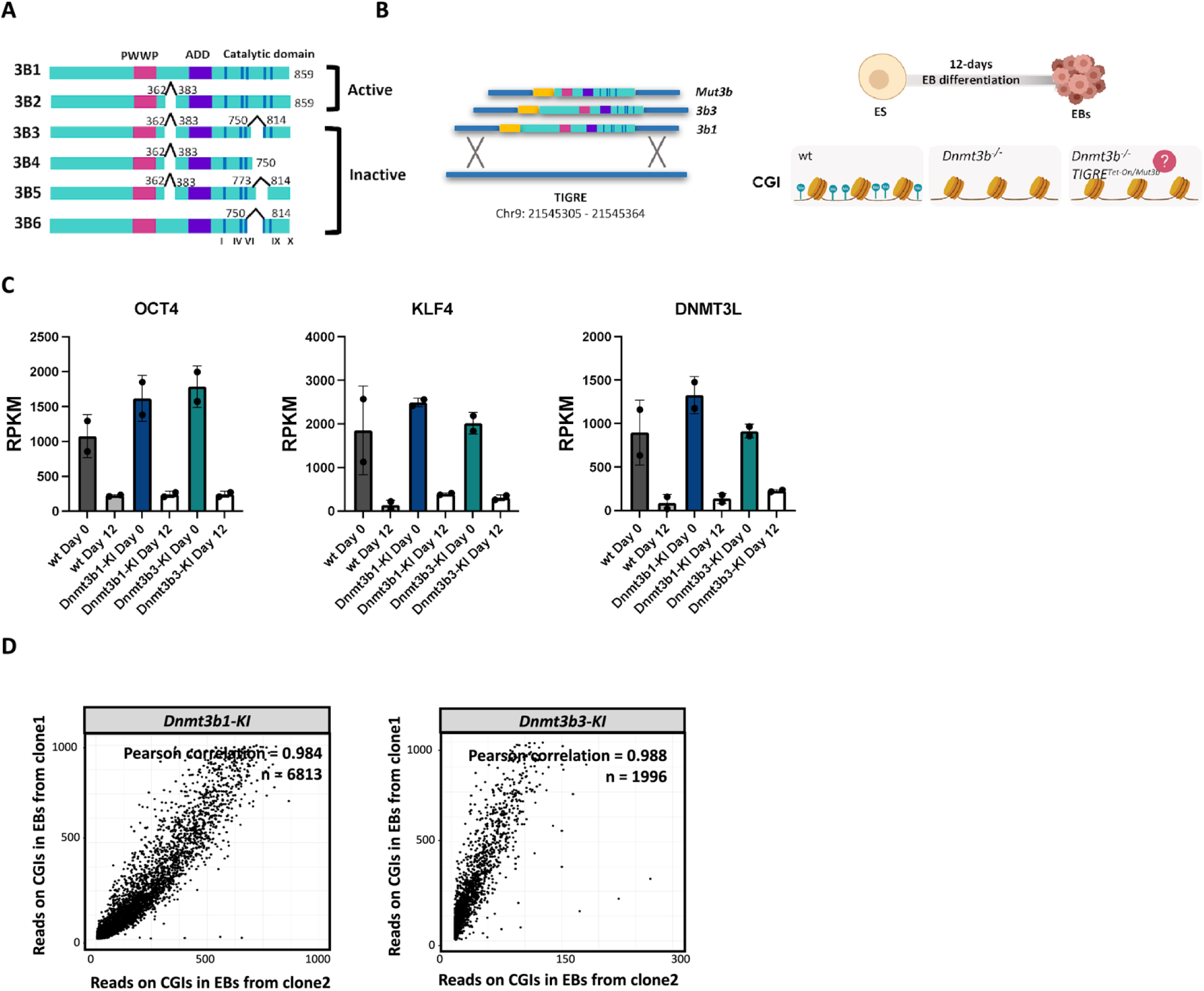
Analysis of Dnmt3b^-/-^ cell lines complemented with Dnmt3b1 or Dnmt3b transgenes. A) Schematic showing isoforms of DNMT3B. DNMT3B1 and 3B2 are both catalytically active. DNMT3B2-3B5 lack amino acids 362-383. The catalytically inactive isoforms of DNMT3B lack residues from the conserved motifs (denoted by the roman numerals on the bottom of the schematic) in the catalytic domain as shown. B) Schematic illustrating the targeting of DNMT3B isoforms or mutants into the TIGRE locus of iXist-ChrX line for generation of complementation lines. The coordinates of the targeted locus are shown at the bottom. C) Expression levels of pluripotency marker genes (*Klf4, Oct4* and *Dnmt3l*) in ES and EBs in wt and DNMT3B1 and DNMT3B3 complementation lines as determined by chrRNA-seq. A trend of downregulation of pluripotency genes can be observed at day 12 indicating differentiation is comparable to wt. A single chrRNA experiment was performed for each of two clones, giving two replicates for the Dnmt3b1-KI cell line. D) Dot plots show correlation between MBD-seq data obtained from 12-day differentiation of two different clones for *Dnmt3b1-KI* and *Dnmt3b3-KI*. Each dot represents a CGI.

**Supplementary Figure 5.**
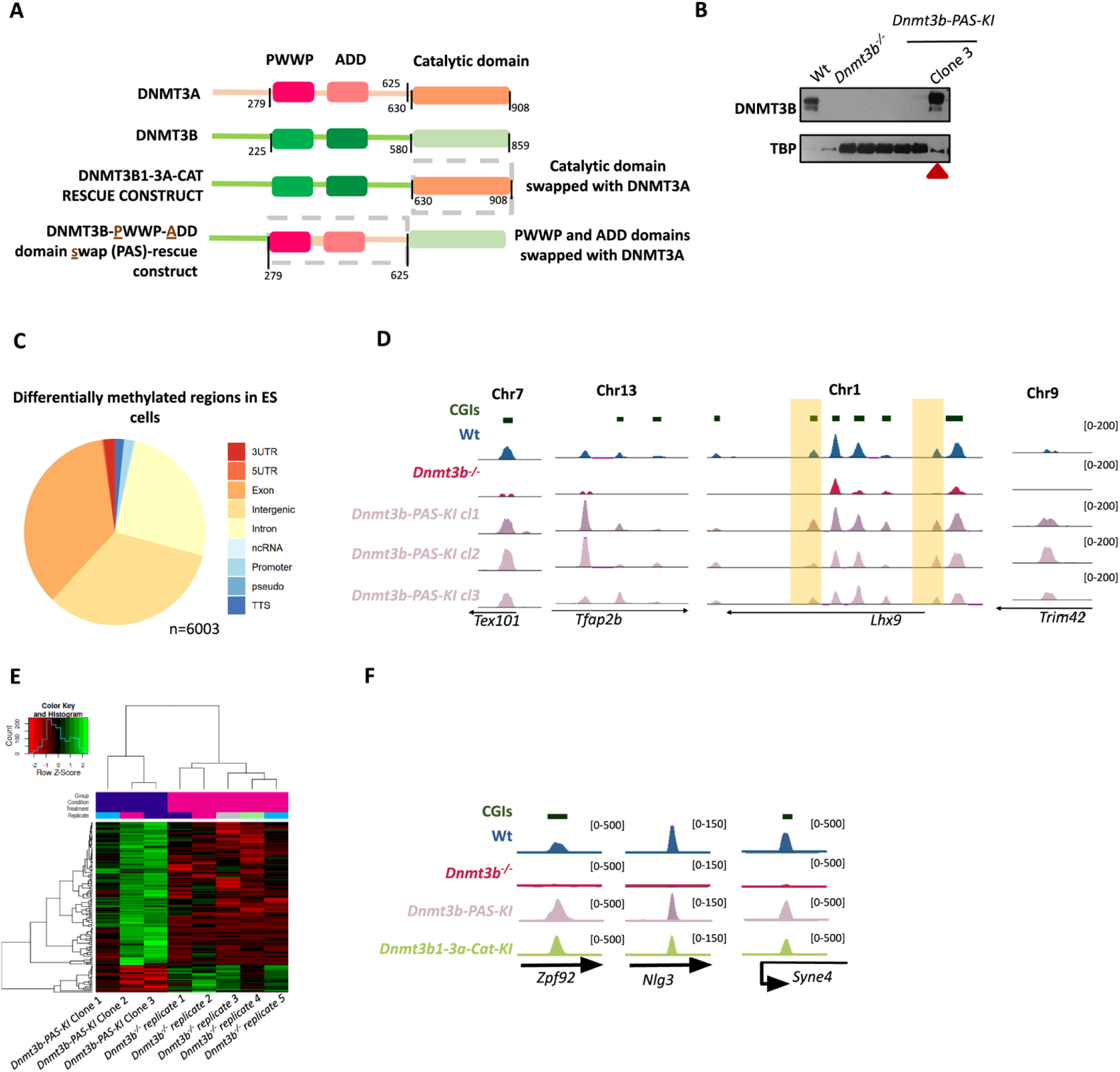
PWWP-ADD of DNMT3B1 is required for Xi CGI methylation. A) Schematic showing chimeric protein construct for *Dnmt3b-3aCat and Dnmt3b-PAS -KI lines.* The DNMT3B catalytic domain (580-859aa) is swapped with that of DNMT3A (630-908 aa). In *Dnmt3b-PAS* the region spanning the PWWP-ADD domain of DNMT3B (225-580aa) is replaced with the equivalent domain of DNMT3A (279-625 aa). B) Western blot showing the expression of DNMT3B-PAS protein in clone3 of *Dnmt3b-PAS-KI* complementation line. C) Pie chart shows the annotation for hyper-methylated regions found in *Dnmt3b-PWWP- ADD_Swap KI* ES cells. The annotation was conducted using homer annotate peaks which uses RefSeq as a reference. D) Genome browser tracks showing selected examples of intergenic MBD-seq signal in *PWWP- ADD_Swap KI* ES cells. E) Heatmap showing the DNMT3B-dependent CpGs methylated in *PWWP-ADD_Swap KI* EBs, recorded as significantly differentially methylated among of 5 replicates for *Dnmt3b^-/-^* and 3 replicates for *PWWP-ADD_Swap KI* EBs. F) Genome browser tracks showing selected examples of 125 DNMT3B-dependent CGIs where methylation was rescued following differentiation of *Dnmt3b_PAS-KI* cells.

**Supplementary Figure 6.**
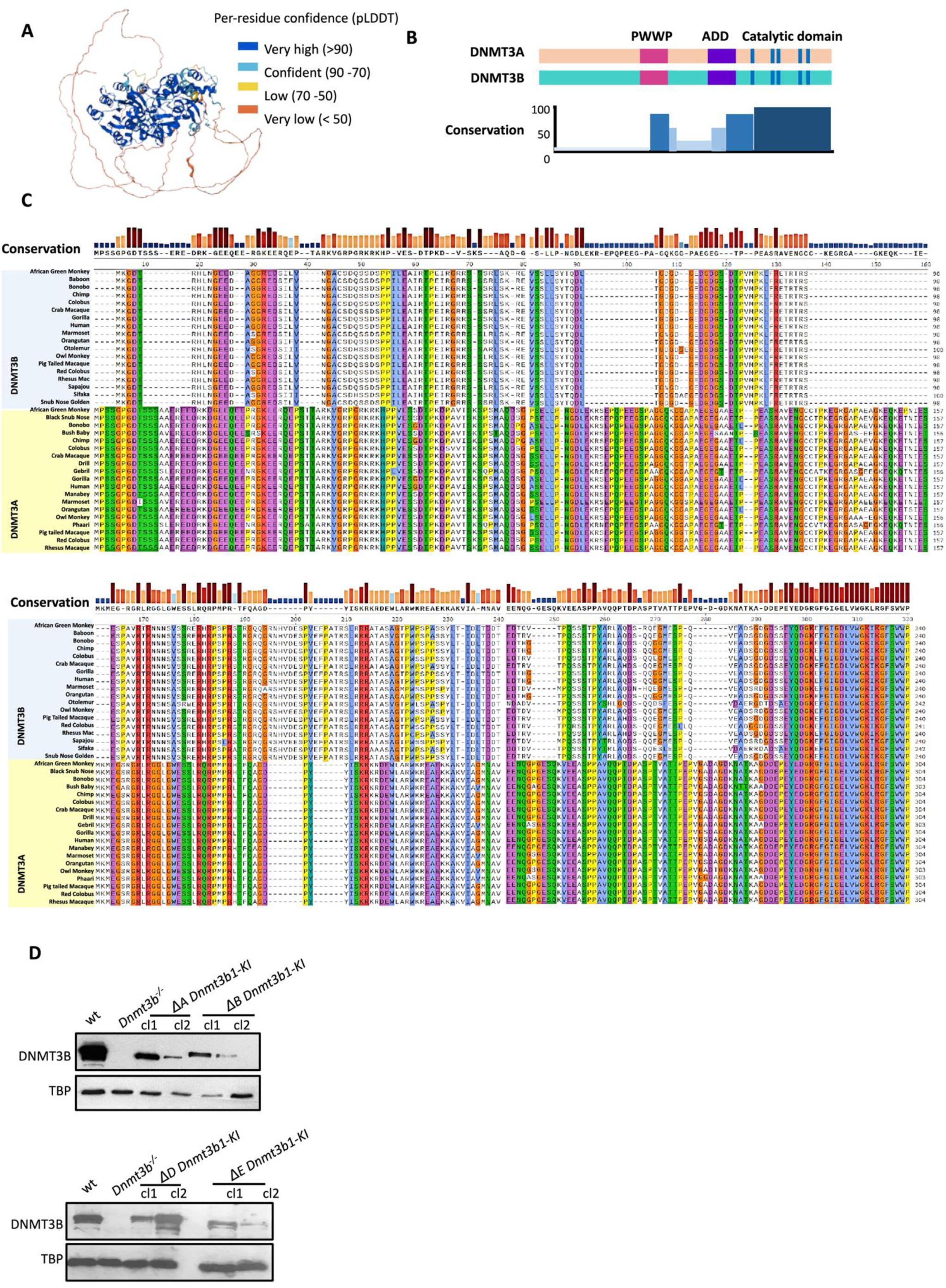
DNMT3B1 N-terminus is required to complement Xi CGI methylation in *Dnmt3b^-/-^* cells. A) Predicted structure of mouse DNMT3B generated by alphafold. Note that the entire N- terminal domain has a very low per-residue confidence score and is shown as an unstructured loop. B) Conservation between mouse DNMT3A and DNMT3B measured by sequence alignment. The conservation score was obtained from Clustal Omega. C) Protein sequence alignment and conservation between DNMT3A and DNMT3B from various mammals. The sequence alignment was performed using Clustal Omega and DNMT3A/B sequences used were as published (Molaro, Malik, & Bourc’his, 2020). D) DNMT3B western blot for clones with various deletions of the N-terminus. TBP loading control.

## References

1. Auclair, G., Guibert, S., Bender, A., & Weber, M. (2014). Ontogeny of CpG island methylation and specificity of DNMT3 methyltransferases during embryonic development in the mouse. Genome biology, 15, 1–16.

2. Baubec, T., Colombo, D. F., Wirbelauer, C., Schmidt, J., Burger, L., Krebs, A. R., … Schübeler, D. (2015). Genomic profiling of DNA methyltransferases reveals a role for DNMT3B in genic methylation. Nature, 520(7546), 243–247.

3. Baylin, S. B., & Jones, P. A. (2016). Epigenetic determinants of cancer. Cold Spring Harbor perspectives in biology, 8(9), a019505.

4. Bird, A., Taggart, M., Frommer, M., Miller, O. J., & Macleod, D. (1985). A fraction of the mouse genome that is derived from islands of nonmethylated, CpG-rich DNA. Cell, 40(1), 91–99.

5. Birke, M., Schreiner, S., García-Cuéllar, M.-P., Mahr, K., Titgemeyer, F., & Slany, R. K. (2002). The MT domain of the proto-oncoprotein MLL binds to CpG-containing DNA and discriminates against methylation. Nucleic acids research, 30(4), 958–965.

6. Blewitt, M. E., Gendrel, A.-V., Pang, Z., Sparrow, D. B., Whitelaw, N., Craig, J. M., … Brockdorff, N. (2008). SmcHD1, containing a structural-maintenance-of-chromosomes hinge domain, has a critical role in X inactivation. Nature genetics, 40(5), 663–669.

7. Borgel, J., Guibert, S., Li, Y., Chiba, H., Schübeler, D., Sasaki, H., … Weber, M. (2010). Targets and dynamics of promoter DNA methylation during early mouse development. Nature genetics, 42(12), 1093–1100.

8. Bourc’his, D. b., Xu, G.-L., Lin, C.-S., Bollman, B., & Bestor, T. H. (2001). Dnmt3L and the establishment of maternal genomic imprints. Science, 294(5551), 2536–2539.

9. Bowness, J. S., Nesterova, T. B., Wei, G., Rodermund, L., Almeida, M., Coker, H., … Brockdorff, N. (2022). Xist-mediated silencing requires additive functions of SPEN and Polycomb together with differentiation-dependent recruitment of SmcHD1. Cell Reports, 39(7).

10. Boyko, K., Arkova, O., Nikolaeva, A., Popov, V. O., Georgiev, P., & Bonchuk, A. (2022). Structure of the DNMT3B ADD domain suggests the absence of a DNMT3A-like autoinhibitory mechanism. Biochemical and Biophysical Research Communications, 619, 124–129.

11. Chedin, F., Lieber, M. R., & Hsieh, C.-L. (2002). The DNA methyltransferase-like protein DNMT3L stimulates de novo methylation by Dnmt3a. Proceedings of the National Academy of Sciences, 99(26), 16916–16921.

12. Chen, Z. X., Mann, J. R., Hsieh, C. L., Riggs, A. D., & Chédin, F. (2005). Physical and functional interactions between the human DNMT3L protein and members of the de novo methyltransferase family. Journal of cellular biochemistry, 95(5), 902–917.

13. Cierpicki, T., Risner, L. E., Grembecka, J., Lukasik, S. M., Popovic, R., Omonkowska, M., … Bushweller, J. H. (2010). Structure of the MLL CXXC domain–DNA complex and its functional role in MLL-AF9 leukemia. Nature structural & molecular biology, 17(1), 62–68.

14. Dahlet, T., Argüeso Lleida, A., Al Adhami, H., Dumas, M., Bender, A., Ngondo, R. P., … Bardet, A. F. (2020). Genome-wide analysis in the mouse embryo reveals the importance of DNA methylation for transcription integrity. Nature communications, 11(1), 3153.

15. Dawlaty, M. M., Breiling, A., Le, T., Raddatz, G., Barrasa, M. I., Cheng, A. W., … Xu, M. (2013). Combined deficiency of Tet1 and Tet2 causes epigenetic abnormalities but is compatible with postnatal development. Developmental cell, 24(3), 310–323.

16. Dobin, A., Davis, C. A., Schlesinger, F., Drenkow, J., Zaleski, C., Jha, S., … Gingeras, T. R. (2013). STAR: ultrafast universal RNA-seq aligner. Bioinformatics, 29(1), 15–21.

17. Duymich, C. E., Charlet, J., Yang, X., Jones, P. A., & Liang, G. (2016). DNMT3B isoforms without catalytic activity stimulate gene body methylation as accessory proteins in somatic cells. Nature communications, 7(1), 11453.

18. Emperle, M., Bangalore, D. M., Adam, S., Kunert, S., Heil, H. S., Heinze, K. G., … Jeltsch, A. (2021). Structural and biochemical insight into the mechanism of dual CpG site binding and methylation by the DNMT3A DNA methyltransferase. Nucleic acids research, 49(14), 8294–8308.

19. Ficz, G., Branco, M. R., Seisenberger, S., Santos, F., Krueger, F., Hore, T. A., … Reik, W. (2011). Dynamic regulation of 5-hydroxymethylcytosine in mouse ES cells and during differentiation. Nature, 473(7347), 398–402.

20. Gao, L., Emperle, M., Guo, Y., Grimm, S. A., Ren, W., Adam, S., … Yin, J. (2020). Comprehensive structure-function characterization of DNMT3B and DNMT3A reveals distinctive de novo DNA methylation mechanisms. Nature communications, 11(1), 3355.

21. Gendrel, A.-V., Apedaile, A., Coker, H., Termanis, A., Zvetkova, I., Godwin, J., … Taylor, S. (2012). Smchd1-dependent and-independent pathways determine developmental dynamics of CpG island methylation on the inactive X chromosome. Developmental cell, 23(2), 265–279.

22. Gordon, C. A., Hartono, S. R., & Chédin, F. (2013). Inactive DNMT3B splice variants modulate de novo DNA methylation. PloS one, 8(7), e69486.

23. Gowher, H., & Jeltsch, A. (2018). Mammalian DNA methyltransferases: new discoveries and open questions. Biochemical Society Transactions, 46(5), 1191–1202.

24. Gowher, H., Liebert, K., Hermann, A., Xu, G., & Jeltsch, A. (2005). Mechanism of stimulation of catalytic activity of Dnmt3A and Dnmt3B DNA-(cytosine-C5)-methyltransferases by Dnmt3L. Journal of Biological Chemistry, 280(14), 13341–13348.

25. Greenberg, M. V., & Bourc’his, D. (2019). The diverse roles of DNA methylation in mammalian development and disease. Nature reviews Molecular cell biology, 20(10), 590–607.

26. Gu, T., Hao, D., Woo, J., Huang, T.-W., Guo, L., Lin, X., … Jeong, M. (2022). The disordered N-terminal domain of DNMT3A recognizes H2AK119ub and is required for postnatal development. Nature genetics, 54(5), 625–636.

27. Guo, X., Wang, L., Li, J., Ding, Z., Xiao, J., Yin, X., … Li, G. (2015). Structural insight into autoinhibition and histone H3-induced activation of DNMT3A. Nature, 517(7536), 640–644.

28. Handa, V., & Jeltsch, A. (2005). Profound flanking sequence preference of Dnmt3a and Dnmt3b mammalian DNA methyltransferases shape the human epigenome. Journal of molecular biology, 348(5), 1103–1112.

29. Hon, G. C., Song, C.-X., Du, T., Jin, F., Selvaraj, S., Lee, A. Y., … Wang, B.-A. (2014). 5mC oxidation by Tet2 modulates enhancer activity and timing of transcriptome reprogramming during differentiation. Molecular cell, 56(2), 286–297.

30. Hu, B., Petela, N., Kurze, A., Chan, K.-L., Chapard, C., & Nasmyth, K. (2015). Biological chromodynamics: a general method for measuring protein occupancy across the genome by calibrating ChIP-seq. Nucleic acids research, 43(20), e132–e132.

31. Ichihara, S., Nagao, K., Sakaguchi, T., Obuse, C., & Sado, T. (2022). SmcHD1 underlies the formation of H3K9me3 blocks on the inactive X chromosome in mice. Development, 149(15), dev200864.

32. Illingworth, R. S., & Bird, A. P. (2009). CpG islands–‘a rough guide’. FEBS letters, 583(11), 1713–1720.

33. Illingworth, R. S., Gruenewald-Schneider, U., Webb, S., Kerr, A. R., James, K. D., Turner, D. J., … Bird, A. P. (2010). Orphan CpG islands identify numerous conserved promoters in the mammalian genome. PLoS genetics, 6(9), e1001134.

34. Ito, S., D’Alessio, A. C., Taranova, O. V., Hong, K., Sowers, L. C., & Zhang, Y. (2010). Role of Tet proteins in 5mC to 5hmC conversion, ES-cell self-renewal and inner cell mass specification. Nature, 466(7310), 1129–1133.

35. Iwase, S., Xiang, B., Ghosh, S., Ren, T., Lewis, P. W., Cochrane, J. C., … Li, H. (2011). ATRX ADD domain links an atypical histone methylation recognition mechanism to human mental-retardation syndrome. Nature structural & molecular biology, 18(7), 769–776.

36. Jeltsch, A., & Jurkowska, R. Z. (2016). Allosteric control of mammalian DNA methyltransferases–a new regulatory paradigm. Nucleic acids research, 44(18), 8556–8575.

37. Jeong, S., Liang, G., Sharma, S., Lin, J. C., Choi, S. H., Han, H., … Jones, P. A. (2009). Selective anchoring of DNA methyltransferases 3A and 3B to nucleosomes containing methylated DNA. Molecular and cellular biology, 29(19), 5366–5376.

38. Krueger, F., & Andrews, S. R. (2016). SNPsplit: Allele-specific splitting of alignments between genomes with known SNP genotypes. F1000Research, 5.

39. Langmead, B., & Salzberg, S. L. (2012). Fast gapped-read alignment with Bowtie 2. Nature methods, 9(4), 357–359.

40. Liao, Y., Smyth, G. K., & Shi, W. (2014). featureCounts: an efficient general purpose program for assigning sequence reads to genomic features. Bioinformatics, 30(7), 923–930.

41. Lin, I. G., Han, L., Taghva, A., O’Brien, L. E., & Hsieh, C.-L. (2002). Murine de novo methyltransferase Dnmt3a demonstrates strand asymmetry and site preference in the methylation of DNA in vitro. Molecular and cellular biology, 22(3), 704–723.

42. Lock, L. F., Takagi, N., & Martin, G. R. (1987). Methylation of the Hprt gene on the inactive X occurs after chromosome inactivation. Cell, 48(1), 39–46.

43. Long, H. K., Blackledge, N. P., & Klose, R. J. (2013). ZF-CxxC domain-containing proteins, CpG islands and the chromatin connection. Biochemical Society Transactions, 41(3), 727–740.

44. Lorsbach, R., Moore, J., Mathew, S., Raimondi, S., Mukatira, S., & Downing, J. (2003). TET1, a member of a novel protein family, is fused to MLL in acute myeloid leukemia containing the t (10; 11)(q22; q23). Leukemia, 17(3), 637–641.

45. Love, M. I., Huber, W., & Anders, S. (2014). Moderated estimation of fold change and dispersion for RNA-seq data with DESeq2. Genome biology, 15, 1–21.

46. Mallona, I., Ilie, I. M., Karemaker, I. D., Butz, S., Manzo, M., Caflisch, A., & Baubec, T. (2021). Flanking sequence preference modulates de novo DNA methylation in the mouse genome. Nucleic acids research, 49(1), 145–157.

47. Mitsuhiro, E., Endo, T. A., Jun, S., Katsuhiko, H., Anca, F., Kit-Wan, M., … Naoko, O. (2017). PCGF6- PRC1 suppresses premature differentiation of mouse embryonic stem cells by regulating germ cell-related genes. eLife, 6.

48. Molaro, A., Malik, H. S., & Bourc’his, D. (2020). Dynamic evolution of de novo DNA methyltransferases in rodent and primate genomes. Molecular biology and evolution, 37(7), 1882–1892.

49. Nesterova, T. B., Wei, G., Coker, H., Pintacuda, G., Bowness, J. S., Zhang, T., … Carter, E. J. (2019). Systematic allelic analysis defines the interplay of key pathways in X chromosome inactivation. Nature communications, 10(1), 3129.

50. Nowialis, P., Lopusna, K., Opavska, J., Haney, S. L., Abraham, A., Sheng, P., … Simpson, M. (2019). Catalytically inactive Dnmt3b rescues mouse embryonic development by accessory and repressive functions. Nature communications, 10(1), 4374.

51. Okano, M., Bell, D. W., Haber, D. A., & Li, E. (1999). DNA methyltransferases Dnmt3a and Dnmt3b are essential for de novo methylation and mammalian development. Cell, 99(3), 247–257.

52. Ono, R., Taki, T., Taketani, T., Taniwaki, M., Kobayashi, H., & Hayashi, Y. (2002). LCX, leukemia-associated protein with a CXXC domain, is fused to MLL in acute myeloid leukemia with trilineage dysplasia having t (10; 11)(q22; q23). Cancer research, 62(14), 4075–4080.

53. Ooi, S. K., Qiu, C., Bernstein, E., Li, K., Jia, D., Yang, Z., … Allis, C. D. (2007). DNMT3L connects unmethylated lysine 4 of histone H3 to de novo methylation of DNA. Nature, 448(7154), 714–717.

54. Pruitt, K. D., Tatusova, T., & Maglott, D. R. (2005). NCBI Reference Sequence (RefSeq): a curated non-redundant sequence database of genomes, transcripts and proteins. Nucleic acids research, 33(suppl_1), D501–D504.

55. Ramírez, F., Dündar, F., Diehl, S., Grüning, B. A., & Manke, T. (2014). deepTools: a flexible platform for exploring deep-sequencing data. Nucleic acids research, 42(W1), W187–W191.

56. Sado, T., Fenner, M. H., Tan, S.-S., Tam, P., Shioda, T., & Li, E. (2000). X inactivation in the mouse embryo deficient for Dnmt1: distinct effect of hypomethylation on imprinted and random X inactivation. Developmental biology, 225(2), 294–303.

57. Schübeler, D. (2015). Function and information content of DNA methylation. Nature, 517(7534), 321–326.

58. Stark, R., & Brown, G. (2011). DiffBind: differential binding analysis of ChIP-Seq peak data. R package version, 100(4.3).

59. Straussman, R., Nejman, D., Roberts, D., Steinfeld, I., Blum, B., Benvenisty, N., … Cedar, H. (2009). Developmental programming of CpG island methylation profiles in the human genome. Nature structural & molecular biology, 16(5), 564–571.

60. Suetake, I., Shinozaki, F., Miyagawa, J., Takeshima, H., & Tajima, S. (2004). DNMT3L stimulates the DNA methylation activity of Dnmt3a and Dnmt3b through a direct interaction. Journal of Biological Chemistry, 279(26), 27816–27823.

61. Taglini, F., Kafetzopoulos, I., Rolls, W., Musialik, K. I., Lee, H. Y., Zhang, Y., … Rubio-Ramon, C. (2024). DNMT3B PWWP mutations cause hypermethylation of heterochromatin. EMBO reports, 25(3), 1130–1155.

62. Weinberg, D. N., Papillon-Cavanagh, S., Chen, H., Yue, Y., Chen, X., Rajagopalan, K. N., … Nikbakht, H. (2019). The histone mark H3K36me2 recruits DNMT3A and shapes the intergenic DNA methylation landscape. Nature, 573(7773), 281–286.

63. Weinberg, D. N., Rosenbaum, P., Chen, X., Barrows, D., Horth, C., Marunde, M. R., … Lu, C. (2021). Two competing mechanisms of DNMT3A recruitment regulate the dynamics of de novo DNA methylation at PRC1-targeted CpG islands. Nature genetics, 53(6), 794–800.

64. Wu, H., D’Alessio, A. C., Ito, S., Xia, K., Wang, Z., Cui, K., … Zhang, Y. (2011). Dual functions of Tet1 in transcriptional regulation in mouse embryonic stem cells. Nature, 473(7347), 389–393.

65. Xu, T.-H., Liu, M., Zhou, X. E., Liang, G., Zhao, G., Xu, H. E., … Jones, P. A. (2020). Structure of nucleosome-bound DNA methyltransferases DNMT3A and DNMT3B. Nature, 586(7827), 151–155.

66. Yagi, M., Kabata, M., Tanaka, A., Ukai, T., Ohta, S., Nakabayashi, K., … Yamamoto, T. (2020). Identification of distinct loci for de novo DNA methylation by DNMT3A and DNMT3B during mammalian development. Nature communications, 11(1), 3199.

67. Zeng, H., Horie, K., Madisen, L., Pavlova, M. N., Gragerova, G., Rohde, A. D., … Kramer, F. (2008). An inducible and reversible mouse genetic rescue system. PLoS genetics, 4(5), e1000069.

68. Zeng, Y., Ren, R., Kaur, G., Hardikar, S., Ying, Z., Babcock, L., … Cheng, X. (2020). The inactive Dnmt3b3 isoform preferentially enhances Dnmt3b-mediated DNA methylation. Genes & development, 34(21-22), 1546–1558.

69. Zhang, Y., Liu, T., Meyer, C. A., Eeckhoute, J., Johnson, D. S., Bernstein, B. E., … Li, W. (2008). Model-based analysis of ChIP-Seq (MACS). Genome biology, 9, 1–9.

